# Primate origins of human event cognition

**DOI:** 10.1101/2023.11.23.568086

**Authors:** Vanessa A. D. Wilson, Sebastian Sauppe, Sarah Brocard, Erik Ringen, Moritz M. Daum, Stephanie Wermelinger, Nianlong Gu, Caroline Andrews, Arrate Isasi-Isasmendi, Balthasar Bickel, Klaus Zuberbühler

## Abstract

Human language relies on a rich cognitive machinery, partially shared with other animals. One key mechanism, decomposing events into causally-linked agent-patient roles, however, has remained elusive with no known animal equivalent. In humans, agent-patient relations in event cognition drive how languages are processed neurally and expressions structured syntactically. We compared visual event tracking between humans and great apes, using stimuli that would elicit causal processing in humans. After accounting for attention to background information, we found similar gaze patterns to agent-patient relations in all species, mostly alternating attention to agents and patients, presumably in order to learn the nature of the event, and occasionally privileging agents under specific conditions. Six-month-old infants, in contrast, did not follow agent-patient relations and attended mostly to background information. We conclude that event role tracking, a cognitive foundation of syntax, evolved long before language but requires time and experience to become ontogenetically available.

**Significance statement:** Human adults and great apes, but not human infants, track natural events as agent-patient relations, a cognitive foundation for syntax.

## Main text

Language is considered unique to humans, a distinction which leads to the prevailing question of how it has evolved. An empirical strategy has been to identify the cognitive mechanisms that language relies on, and to reconstruct their evolutionary history using comparative research with humans and other animals. One important cognitive mechanism is the propensity for speakers and listeners to decompose events into causally structured agent-patient relations (1). For example, a sentence like “Alice picked up the caterpillar” has Alice as the agent and the caterpillar as the patient. This distinction is deeply entrenched in meaning, neuroanatomically detectable (2) and responsible for core syntactic phenomena, such as case marking or constituent hierarchies (3), with only very few exceptions across the world’s languages (4). Furthermore, languages privilege agent over patient roles, preferring the simplest and default expressions for agents (5), even though this often incurs uncertainty and additional neural activity during sentence planning (6,7). Correspondingly, agents tend to be named before patients (8,9), a trend only matched in sentence structure by a concurrent preference for placing reference to humans before reference to inanimate things (10).

These biases in linguistic expression build on resilient mechanisms in human event cognition (11). For instance, when apprehending the gist of events from still pictures, people tend to be quicker to identify agents than patients (12), and assign agency almost instantly and with remarkably little variation across cultures and languages (13). Early agent identification is typically followed by distributed attention between agents and patients in a processing stage known as ‘relational encoding’ during early sentence planning (6). The same resilience is also apparent in comprehension during sentence processing. When sentences violate expectations about agent and patient roles, for example, by putting patients before agents, neurophysiological measures indicate that agency is usually assigned as the initial default, even when this goes against usage probabilities and the rules of grammar (5,14,15). Together, these findings suggest that language builds on a universal neurocognitive mechanism of event decomposition to make sense of the world and its linguistic representations.

This raises the question of how human event cognition has evolved. We are not aware of any evidence in the animal communication literature that demonstrates that signals can refer to agent-patient interactions, neither in natural communication nor with artificial languages (16). One hypothesis, therefore, is that nonhuman animals (from here on, ‘animals’) do not possess the cognitive resources for decomposing events into agents and patients. Certainly, animals can comprehend aspects of physical causality (e.g., that pushing causes falling) (17). Still, it is unclear whether this is due to perceptions of simple co-occurrences or more complex perceptions of events as agent activities causing patient changes. Related to this, although there is little doubt that animals perceive the participants of events, their attention may be absorbed by the participants’ social attributes, such as their identities, social roles (18), or behavioral intentions (17,19), all of which predict large situational and individual variation in how events are processed.

The alternative hypothesis, to be tested here, is that animals are capable of human-like event decomposition (1), but do not have the motivation or the resources to communicate about agent-patient relations. To explore this, we tested how participants across closely related species of hominids perceived a range of naturalistic events that would elicit causal processing in humans. We compared gaze responses to short video clips between members of the four genera of great apes - humans (*Homo sapiens*), chimpanzees (*Pan troglodytes*), gorillas (*Gorilla gorilla*) and orangutans (*Pongo abelii*). We also tested human infants at 6 months old, before they start to actively use language and whilst still developing linguistic processing abilities. By this age, infants already show an impressive cognitive toolkit: they are sensitive to goal-directed actions and agency (20), track changes in goal-directed behavior (21), and extract key information from video stimuli to understand events (22). At the same time, young infants struggle to process goal predictions (23) and third-party social interactions (24). As infants develop, language and event perception become increasingly intertwined, as documented by the way verbs and actions are processed (25).

Currently, our understanding of agent-privileged event cognition in humans rests mainly on paradigms that use static stimuli, which are often artificial or overly simplistic, and do not reflect the complexity of real-life interactions. In this study, we used dynamic scenarios across a broad range of natural events to compare overt visual attention to agents and patients as the actions unfold. Scenarios were presented as *N*=84 short (2-10 s long), silent video clips, depicting animate agents and both animate and inanimate patients of (unfamiliar) humans, chimpanzees, gorillas, and orangutans engaged in natural interactions. All participants saw the same stimuli (Supplementary Material Table 1). When creating the videos, we deliberately avoided rigidly controlling for low-level perceptual features, as this would have created sterile footage with low socio-ecological validity (26) and, critically, reduced interest for ape participants. Instead, we presented scenes that sought to capture much greater variation of real life. Possible confounding factors, such as differences in the amount of agent motion or relatively larger sizes of agents or patients between videos, were accounted for in the statistical models (see Methods section in the Supplementary Material; Supplementary Material Figures 5 & 6).

For human adults, we expected to see early and overall agent biases, consistent with previous findings, but with attention patterns mediated by the progression of the action rather than the need to extract agent-patient information rapidly, as in brief exposure studies (12). We predicted that if event decomposition were a general feature of great ape cognition and present without language, then visual attention should not differ across the four species. In particular, we predicted earlier attendance to agents than patients, in line with the privileged status of agents in language and gist apprehension in still pictures. Alternatively, if event decomposition were uniquely human — or dependent on language — we expected to find this pattern only in adult humans, and large and random variation in the nonhuman primates, which would likely depend on low-level features such as color or contrast of the stimuli. Regarding the infants, if event-role decomposition required experience gained through observing third-party interactions, we expected to see differences in how they attended to agents and patients compared with adults.

For the analysis, we ran two types of models. Firstly, we conducted a Bayesian time series analysis (for details, see *Statistical analyses* in the methods), for which we split the videos into scenes depicting social interactions (i.e., with animate patients) and scenes depicting interactions with inanimate patients. Preliminary examination of the data revealed large heterogeneity in the time course data between videos, which was not captured by either differences in inanimate and social videos, or relative AOI size (see Methods for more details on accounting for differences in AOI size). To determine where these gaze differences came from, we explored the videos with heterogeneous responses, which concerned specifically inanimate videos of apes. From these scenarios, we identified four categories that might explain the resulting variance: 1) Scenes generally containing food, 2) tool use, such as nut cracking or honey dipping, 3) instances where the agent or patient were oriented towards the camera, and 4) instances where the agent or patient had direct gaze towards the camera, something that could have been perceived as threatening by some ape participants. To estimate the effects of these categories, we fitted additional models (figures for these models can be viewed in a dashboard within the Supplementary data, at https://osf.io/47wap/?view_only=8c2b20667fc441178269291fda5262bf). These models indicated a clear difference in gaze patterns to inanimate scenes containing food, compared to those containing objects. We thus further split the inanimate videos into those containing food or not.

GLMMs such as ours are best understood by examining model predictions given different values of the covariates (time, species, condition), rather than interpreting individual coefficients. This is due to both the large number of parameters in multilevel models and the non-linearity induced by the link function. As such, we draw on the credible intervals of the models’ predictions at multiple time points to interpret the data (27) (see, for example, Figures 1 and 2). Four main findings emerged. Firstly, in both adult humans and apes, after accounting for attention to background information, there was an early focus on the action with attentional switches between agents and patients, suggesting participants engaged in relational encoding, similar to what humans do when asked to describe actions depicted in still images (Figure 1; Supplementary Material Table 3). Secondly, in both adult humans and apes, the inanimate food condition triggered a striking bias towards agents at the onset of the action (with posterior probabilities for log odds ratios of agent vs. patient fixations all being above 0), absent from the other conditions (Figure 1). For humans, agents remained salient throughout the scenario, whilst for apes, this gaze pattern was less pronounced. Thirdly, human adults directed most of their visual attention to agents and patients, whilst apes attended more to other information. This effect was slightly stronger for orangutans, suggesting a possible phylogenetic emergence of event role attribution (Figure 2; Supplementary Material Tables 4 - 5; Supplementary Material Figures 2-4). Finally, human infants radically differed from the other groups, by attending mostly to other, non-agent and non-patient, information within each scene (Figure 2).

**Figure 1:**
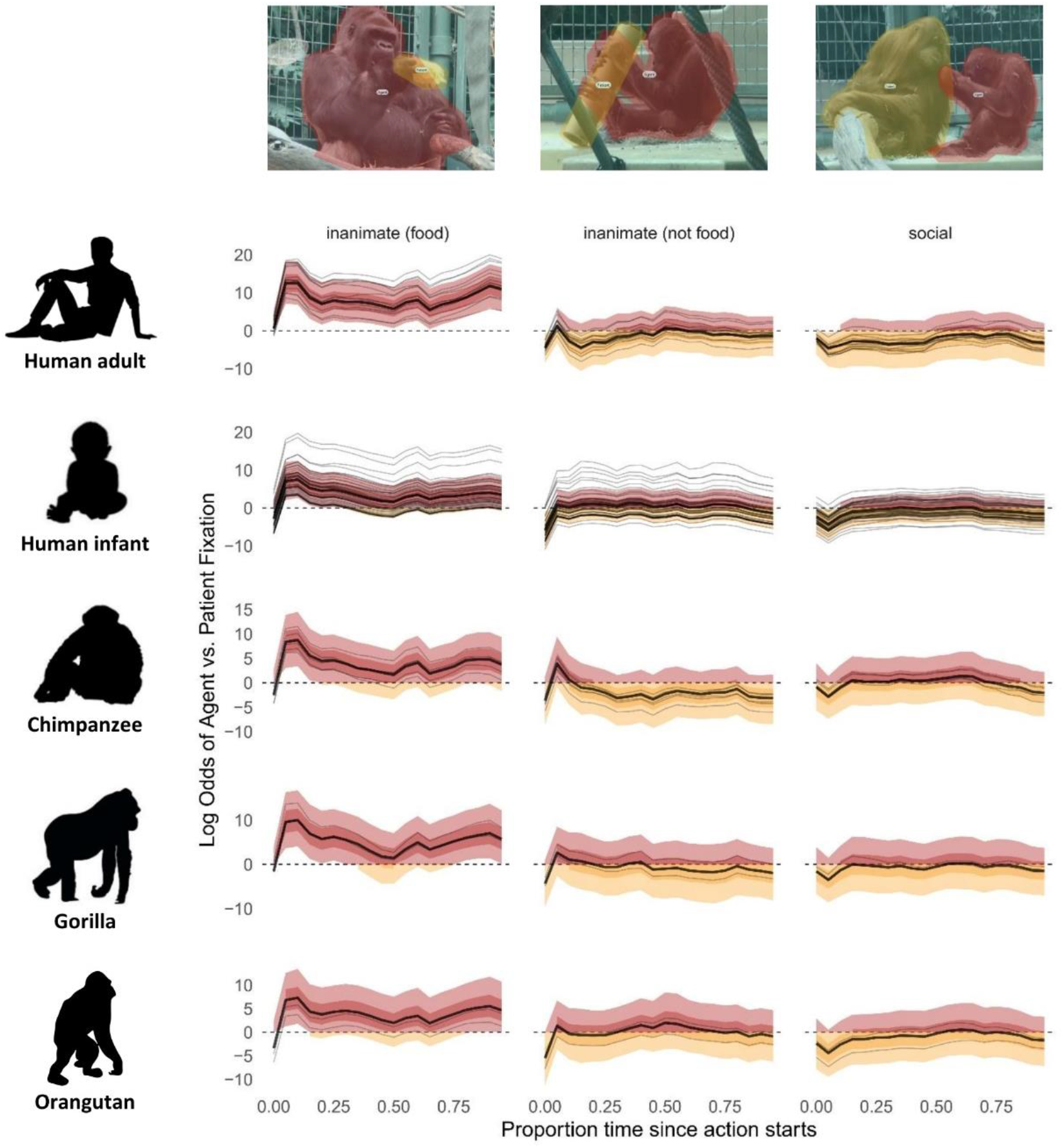
Model predictions of log odds ratio for fixation to either agent (red) or patient (orange) over time. Thick lines represent the grand mean, thin lines represent individual participants. Time point 0 on the x-axis indicates action start time, normalized across stimuli. Light shaded ribbons indicate 90% credible intervals. When these exceed 0, there is a 90% probability of gaze to the agent (light red); when they are below 0, there is a 90% probability of gaze to the patient (light orange); when they include zero, there is a 90% probability that gaze alternates between agent and patient. The darker shaded ribbons indicate the middle 50% of the posterior probability mass.

**Figure 2:**
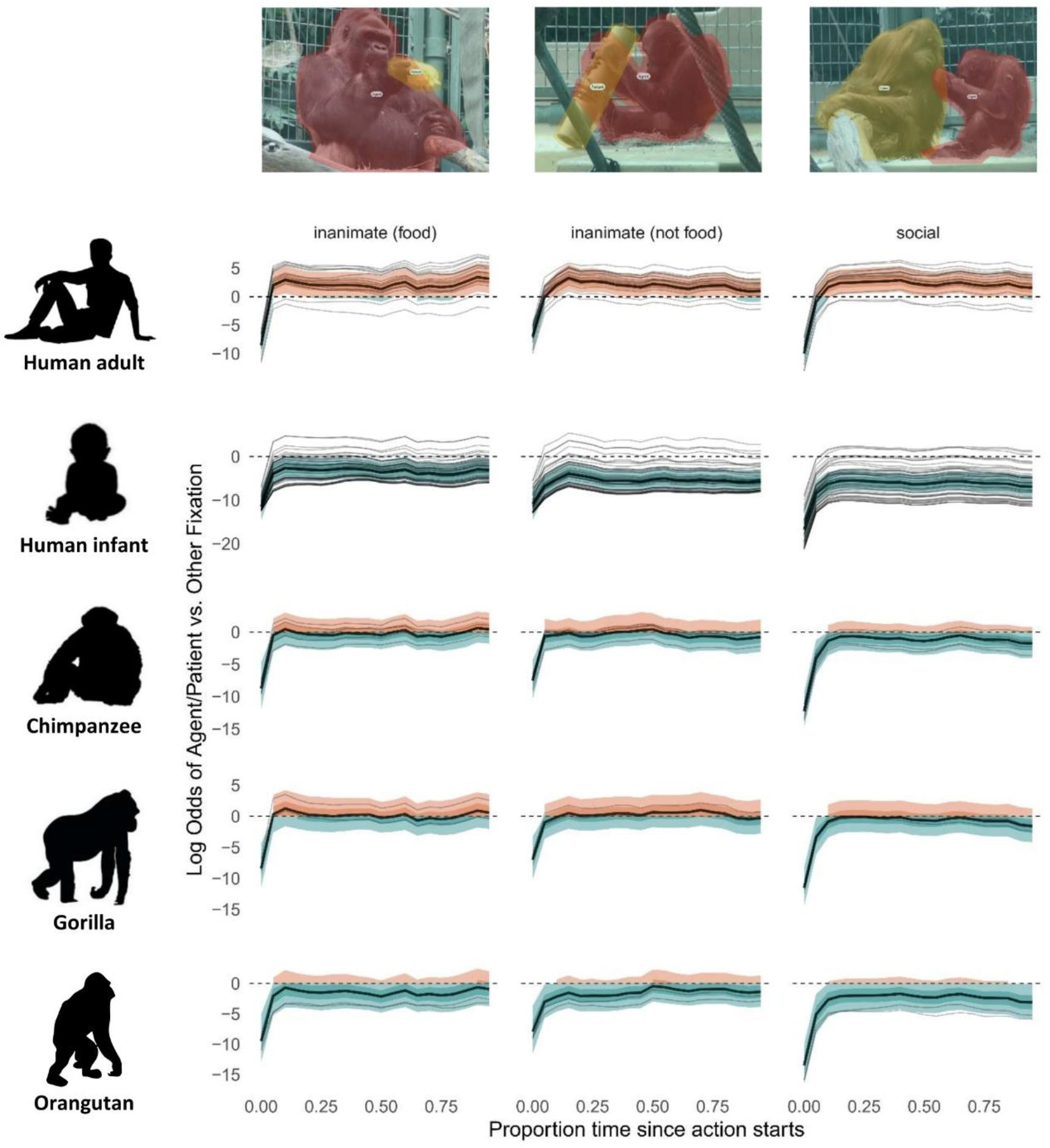
Model predictions of log odds ratio for fixation to agent or patient (orange) versus other (turquoise) over time. Thick lines represent the grand mean, thin lines represent individual participants. Time point 0 on the x-axis indicates action start time, normalized across stimuli. Light shaded ribbons refer to 90% credible intervals. They exceed 0 for humans, indicating gaze priority for agents and patients (orange). They are below 0 for human infants (turquoise) (with the exception of inanimate food), indicating gaze primarily to background information. 90% credible intervals for apes border around 0, indicating that attention is divided between agents and patients (orange) and other information (turquoise). Darker shaded intervals indicate the middle 50% of the posterior probability mass.

In addition to the time series analysis, we fitted a Dirichlet model that examined whether accounting for event roles better explained gaze patterns than any of the covariates (such as relative size of agents and patients), by comparing two models that differed in their expectations about gaze proportion to agents and patients (for details, see *Trial Averaged Model Comparison* in the methods). A null model predicted that, assuming event roles were not needed to differentiate two entities, gaze proportion to agents and patients should be equal. An alternative model predicted that, assuming event roles better explained gaze patterns to two entities, gaze proportion would differ between agents and patients. Results indicated that the non-null model (i.e., assuming different gaze proportions to agents and patients) better predicted the gaze patterns, indicating that participants do not simply differentiate between two entities, but between the roles attached to those entities (Table 1).

**Table 1.** Model comparison differentiating event role specification as a predictor of agent-patient gaze proportion

| Human adults |  |  |  |  |  |
| --- | --- | --- | --- | --- | --- |
| | ELPD<sub>LOO</sub> | SE(ELPD<sub>LOO</sub>) | ΔELPD | SE(ΔELPD) | weight |
| *Null model* | 1047.6 | 39 | -509.9 | 43.5 | 0.058 |
| *Alternative model* | 1557.5 | 46.2 |  |  | 0.942 |
| Human infants |  |  |  |  |  |
| | ELPD<sub>LOO</sub> | SE(ELPD<sub>LOO</sub>) | ΔELPD | SE(ΔELPD) | weight |
| *Null model* | 8762.4 | 184.4 | -191 | 22.3 | 0.086 |
| *Alternative model* | 8953.3 | 181.4 |  |  | 0.914 |
| Apes |  |  |  |  |  |
| | ELPD<sub>LOO</sub> | SE(ELPD<sub>LOO</sub>) | ΔELPD | SE(ΔELPD) | weight |
| *Null model* | 1325.9 | 63.8 | -167 | 23.2 | 0.086 |
| *Alternative model* | 1492.9 | 65.6 |  |  | 0.914 |

Note. *Null model* = without event role as predictor. *Alternative model* = with event role as predictor. ELPD<sub>LOO</sub> is the expected log pointwise density, which indicates the predictive accuracy of the model. $\Delta$ ELPD indicates the difference in predictive accuracy between the null model and the alternative model. Model weights indicate the probability that the alternative model will predict new data better than the null model.

Evolutionary theories of syntax have focused mainly on how formal complexity has emerged (28–30), whereas the underlying cognitive mechanisms have rarely been addressed. Here, we tested a cognitive hypothesis, which proposes that central aspects of human syntax, such as case-marking or constituent hierarchy, build on a prelinguistic cognitive mechanism that decomposes events into causally structured agent-patient relations (1). To test this, we exposed apes to stimuli that elicit causal processing in humans, and compared the gaze patterns between humans and nonhuman apes. Participants across species tracked events in strikingly similar ways, focusing on the action between agents and patients in a manner reminiscent of relational encoding for planning to speak (6). This finding suggests that apes, like humans, decompose the causal agent-patient roles depicted. The only noticeable difference was that nonhuman apes showed more visual exploration of background information than humans, perhaps due to differences in experience with watching and interpreting videos or higher intrinsic interest in scanning the larger environment. This is reflected in Supplementary Figures 2 and 3, where human adult attention to agents and patients is more pronounced, because these results do not account for attention to ‘other’ information. Notably, apes’ looking behavior showed more similarity to human adults than did human infants. If apes were unable to track agent-patient relations, we would expect attention patterns similar to those seen in infants. These observations reject the hypothesis that event decomposition emerged as a unique form of human cognition together with language.

Unexpectedly, across all event categories, in neither humans nor great apes did we find a strong bias towards agents (with the exception of food scenarios). This is in contrast to findings from a large body of previous research using static images, as well as a recent comparative study examining event role preferences (31). This difference is likely due to the nature of the stimuli. Static stimuli tap into rapid event perception (32) and usually capture the midpoint of an event, which increases the difficulty in identifying distinct roles, similar to when listeners have to come up with rapid predictions on roles while processing a sentence (33,34). It is likely that an agent bias manifests itself primarily under these high-demand conditions, while it is not as relevant when watching an event that unfolds over time (35).

Additionally, unlike some previous studies, we controlled for size and movement of the areas of interest, as well as event type; size of the agent or patient, in particular, has a strong effect on gaze probability (Supplementary Material Figure 5). Curiously, when considering gaze differences between event categories, we found the strongest agent bias in video scenes depicting interactions with food. One possible explanation is that, in social species, attending to agents who have access to food provides a survival advantage. This points to social learning as a possible precursor for semantic role attribution. An intriguing possibility requiring further studies, therefore, is that the agent bias reported elsewhere has its roots in trophic interactions. Further research is needed to more systematically explore different degrees and kinds of cognitive pressure in event cognition and different ecological contexts of events.

The 6-month-old infants attended to agents and patients with very low probability. Perhaps this was due to a lack of experience with videos, which is somewhat inconsistent with the fact they showed an initial agency bias in food scenarios, and the fact that young infants have been able to attribute agency in other experiments (36). A more plausible explanation may be that infants this young have trouble detecting causality in complex visual material (37), especially in social interactions. Indeed, our stimuli were more complex than typically used in infant studies. However, these were chosen to reflect real-life diversity of events. Key ingredients, such as event parsing (38), causal integration across scenes (23) and triadic awareness (39), are known to develop gradually during the first 12 months, suggesting that our content was too challenging and probably too alien to their existing world experience. Another plausible hypothesis is that processing of dynamic natural scenes requires computational resources and oculomotor control not yet sufficiently developed at this age (40). As a consequence, integration and interpretation of relevant information is more time-consuming (22) or just not yet possible. Given that event categorisation relies on neurally constructed models which are updated with experience (41), it is likely that at this age, infants are still developing the event models that will guide their attention to scene information.

In sum, our study demonstrates that nonhuman great apes share with human adults the cognitive mechanism required for tracking agent-patient relations, which naturally triggers the question of why they do not have language. Earlier explanations that relied on human-animal morphological differences in vocal tracts (42), lack of declarative communication (43), and lack of call composition (44) no longer stand. Our results add to the shared cognitive foundations of language by indicating that event decomposition, a foundation of syntax, evolved before language, on par with signal combinations (45), theory of mind (46) and joint commitment (47).

What has happened then during human evolutionary history that allowed us to map event roles into verbal expression? We can think of three probably interlinked evolutionary transitions that may have paved the way from primate-like communication to human language: (a) changes in social cognition, (b) changes in communicative needs and (c) changes in expressive power. Regarding social cognition, the key step may have been to externalize event cognition through language, by moving from implicit to explicit attributions. For example, compared to chimpanzees, adult humans attend more to an agent’s face following an unexpected action outcome, as if explicitly trying to discern the actor’s mental state (48). Regarding communicative needs, one argument is that increased levels of cooperation and coordination brought about increased communicative needs, a convergent evolution independent of wider cognitive evolution (49,50). Finally, regarding expressive power, modern humans roughly have three-fold larger brains than chimpanzees, with vastly more computational power, allowing for processing of more varied signal structures. Although there are examples of limited compositionality in animal communication, there is no evidence for free variation and creative use (45). Testing these hypotheses may provide further answers in the quest for the origins of language by better understanding why nonhuman apes do not communicate in the same way as humans do, despite an increasingly closing gap with human cognitive abilities.

## Acknowledgements

For support and assistance at Basel Zoo we thank Adrian Baumeyer, Fabia Wyss, Raphaela Heesen, Stephan Lopez, Gaby Rindlisbacher, Rene Buob, Roland Kleger, Jonas Schaub, Nicole Fischer, Reto Lehmann, Corinne Zollinger, Markus Beutler, Dominic Hohler, Patrick Wyser, Flurin Baer, Amanda Spillmann, Stephan Argast, and the technician team. We also thank Carla Pascual for help with data collection. For providing footage from apes, we thank Emily Genty, Cat Hobaiter, Jennifer Botting, Erin Stromberg, and Atlanta Zoo, as well as Zurich Zoo for allowing us to film their apes. We thank Sara I. Fabrikant and Tumasch Reichenbacher for providing their eye-tracking laboratory for the human adult data collection and Sina Nägelin, Nina Philipp, and Deborah Lamm for collecting the data from human adults and infants, as well as Marco Bleiker for technical assistance with infant data collection. We thank Sebastien Quigley and Carla Pascual for assistance in data processing. We also thank Shreejata Gupta, Christopher Krupenye, and Josep Call for their advice on establishing the eye-tracking setups for data collection from great apes.

## Funding

Swiss National Science Foundation (project grant numbers 310030_185324, K.Z., and 100015_182845, B.B.)

The National Center for Competence in Research “Evolving Language” (SNSF agreement number 51NF40_180888, B.B., K.Z., M.M.D., and Top-Up grant number N603-18-01, V.A.D.W., K.Z., B.B., M.M.D.)

Foundation for Research in Science and the Humanities at the University of Zurich (grant number 20-014, V.A.D.W., S.S., B.B.)

Seed money grant, University Research Priority Program “Evolution in Action”, University of Zurich (S.S.).

## Author contributions

Conceptualization: VADW, SS, SB, MMD, SW, BB, KZ

Methodology: VADW, SS, SB, MMD, SW, BB, KZ

Investigation: VADW, SB Software: SS, ER, NG Validation: ER

Formal analysis: ER, VADW, SS, BB, KZ

Resources: VADW, SS, SB, MMD, CA, AI, BB, KZ

Data curation: VADW, SS, SB, SW, ER, NG

Visualization: ER, VADW, SS, BB

Writing - original draft: VADW

Writing - review and editing: VADW, SS, SB, MMD, EK, SW, NG, CA, AI, BB, KZ

Supervision: BB, KZ

Project administration: VADW

Funding acquisition: BB, KZ, VADW, SS, MMD

**Competing interests:** The authors declare that they have no competing interests.

**Data and Materials availability:** Data and code are available at https://osf.io/47wap/?view_only=8c2b20667fc441178269291fda5262bf

## Supplementary Materials

### Methods

#### Participants

##### Nonhuman apes

Nine great apes from three species were tested. These included five chimpanzees (3 females, age in years: mean = 6.4, SD = 6.14, span = [2.5, 17]), two gorillas (1 female, both 5 years old), and two orangutans (1 female, 15 and 19 years old). The youngest participants included already showed some level of independence from their mothers and interacted freely with their peers. All participants were housed in social groups at Basel Zoo, Switzerland (chimpanzees: 8 adults [2 male], 4 juveniles [2-6 years; 1 male], 1 infant [< 1 year]; gorillas: 5 adults [1 male], 2 juveniles [5 years old; 1 male], 1 infant [< 1 year old]; orangutans: 6 adults [3 male], 3 juveniles [2-7 years; 1 male]). Each group had access to both an indoor and an outdoor enclosure (chimpanzees, indoor: 1096.93 m^3^, outside: 3860.1 m^3^; gorillas, indoor: 1148.84 m^3^, outside: 3025.05 m^3^; orangutans, indoor: 1726.69 m^3^, outside: 8965.1 m^3^).

Ethical approval for the ape research was provided by the Canton of Basel Veterinary Office (approval numbers 2983 and 3077) and by the Animal Welfare Officer at Basel Zoo. All apes participated voluntarily. They were not separated from their group during testing, nor were they food or water deprived, and could leave at any time. They were rewarded for participation with diluted sugar-free syrup, provided in restricted quantities as approved by the zoo’s veterinary team.

##### Humans

Fourteen human adults (7 female, age in years: mean = 26.79, SD = 6.73, span = [18, 41]), recruited among the students of the University of Zurich, participated for payment. Twenty-nine human infants (16 female, age in days: mean = 191.79, SD = 11.80, span = [168, 219]) were recruited through the research unit Developmental Psychology: Infancy and Childhood at the University of Zurich. Infants of parents who speak a language with an ergative case system (e.g., Basque or Hindi) were excluded from recruitment because ergative case is known to modulate agent attention (13). Consistent with general drop-out rates for infant studies (51), data from 15 additional infants were excluded from analyses because they had overall poor or missing calibrations, were not attending to the stimuli, or because (in one case) the mother spoke a language with an ergative case marking system.

All procedures were approved by the local ethics committee of the University of Zurich (approval numbers 18.10.9 and 21.9.18) and performed in accordance with the ethical standards of the 1964 Helsinki Declaration and its later amendments. All adult participants or the infants’ caregivers gave informed consent before data collection. After completing the task, adult participants received CHF 20 and infant participants were rewarded with a certificate and a small present worth approximately CHF 5.

#### Materials

Eighty-four video clips were selected as stimuli to be presented during testing, including 52scenes of apes engaged in natural interactions (see Supplementary for details) and 32 scenes of humans performing various actions. The stimuli depicted either interactions between two animate participants (apes interacting with other apes of the same species or humans interacting with other humans; chimpanzees: 7, gorillas: 6, orangutans: 7, humans: 16) or apes or human interacting with an inanimate object (chimpanzees: 9, gorillas: 9, orangutans: 14, humans: 16). All stimulus videos were between 2 and 10 seconds long (mean = 6.37, SD = 2.08). Animate videos with humans depicted dyadic actions such as *kick*, *pull*, *brush*, *beckon*, *scold*, or *guide*; inanimate videos with humans depicted actions such as *opening a box*, *pouring water*, *playing the drums*, *reading a book, fixing a bicycle*, or *putting on gloves*. We selected actions that we judged the apes to find generally familiar and avoided any actions that apes might find alarming or induce strong emotional arousal (52). A full list of stimuli shown is available in the Supplementary Material Table 1.

Only same-sex pairings were shown in interactions between apes or humans to avoid that the ape participants inferred potential dominance differences between males and females (53), which could affect gaze patterns. Stimuli were presented at 24 frames/second and without sound. All stimuli were edited in Lightworks (version 1.0). For each stimulus video, a mirror-flipped version was generated so that the agent could be presented on both the left and right of the screen. We counterbalanced the agent’s side for each stimulus between participants.

Footage of humans was filmed for the purpose of another study on human event cognition (13). Human actors wore colored shirts; we distributed the colors of agent and patient shirts across scenarios as evenly as possible to account for the possibility that the apes would respond to the color as a cue, rather than attending to the agent/patient dynamic (an overview of shirt color distributions is given in Supplementary Material Table 1).

#### Apparatus and Procedure

Video stimuli were presented at a resolution of 1920*1080 pixels in the center of the 23-inch screen which was positioned above a Tobii TX300 eye tracker (for all participants but one orangutan) or a 23.8 inch screen positioned above a Tobii Spectrum eye tracker (for one orangutan; data for this participant was collected at a later date when a more optimal ape calibration option was available through the Spectrum), while the same device also recorded participants’ gaze (300 Hz sampling rate). The eye tracker was controlled by a Notebook Captiva NH50_70RA laptop computer using the Tobii Pro Lab software.

Each trial started with an image of a piece of fruit serving as a “fixation cross”, placed at either the top or bottom of the screen (always in the same position, with presentation point counterbalanced between trials), so that participants did not already look at the position of the video stimulus when it started, since depicted actors may be visually and conceptually more salient if they happen to be looked at from the outset of stimulus presentation. Human adults, for example, are more likely to choose an event participant as the syntactic subject when already looking at the position where this event participant is located when a stimulus appears (leading to more passive sentence responses during picture description when patients are looked at first) (54–57). Ensuring that participants always fixated outside of the video stimulus boundaries guards against such inadvertent salience effects. Once participants fixated the fruit, the experimenter initiated the video stimulus, lasting between 2 and 10 seconds (see Supplementary Material for example trials).

##### Apes

Two types of setups were used for testing. For gorillas and chimpanzees, the eye tracker and monitor were positioned in a small enclosure (gorillas: floor: 160 * 135 * 165 * 40cm, height: 3 m; chimpanzees: floor: 116 * 91.5 * 160 * 93.5cm, height: 267 cm), separated from the main indoor enclosure by a sliding door and a plexiglass window (55 * 80 cm). Equipment was accessed through a separate door that opened into the keepers’ corridor (see Supplementary Material Figure 1A–B). At the start of each session, the door was raised and the apes could approach the testing area and sit in front of the screen. A fixed drinking nozzle was fitted in the center of the window, providing apes with access to diluted syrup. Provisioning of the syrup was controlled through a tube and catheter.

For the orangutans, the enclosure design made it difficult to implement the use of one fixed setup accessible to all participants. Instead, we implemented a height-adjustable mobile wagon (200.4±50 * 102 * 70 cm) (see Supplementary Material Figure 1, C) to present the eye tracker to participants through the enclosure mesh. The eye tracker was protected by a plexiglass box (63 * 102 * 70 cm) and as with the fixed setups, had a drinking nozzle on the front panel that could be accessed through the mesh. The drinking nozzle was designed to be positioned on a vertical mesh bar, so the apes’ eyes were positioned between the bars.

Sessions took place Monday-Friday in morning (09:00-12:00 hrs) and afternoon sessions (13:30-15:30 hrs). Prior to testing with the apes, we presented a series of training clips of varying stimuli (see Supplementary text) to ensure that they did not find certain types of video stimuli aversive, and we monitored their responses closely. During participation, apes could drink the syrup freely, and could choose whether or not to look at the screen. Overall, those apes who chose to participate in the study showed high motivation for viewing the stimuli.

Once habituated to the apparatus, the eye tracker was calibrated to the apes using a 2-point calibration procedure by displaying a short video in the corners of the screen. Calibration was recorded once, prior to all testing. Calibrations were accepted if tracking accuracy and precision were within 1° visual angle.

Each session began with a green screen for 2 seconds that indicated the start of testing. This was followed by a 4-point validation procedure to check the calibration accuracy against four fixed points on-screen. Following this, a ‘practice’ video was shown, which was not used for data collection but was designed to engage the viewer prior to the presentation of the stimuli of interest. The practice video either depicted humans or apes, so that the content of the videos was predictive of whether participants would see humans or apes in the following trials.

Trials were presented sequentially, as long as the ape kept attending to the screen and did not display any signs of distraction. All stimuli with apes as agents were presented first, followed by all stimuli with human agents. In case a participant became distracted and did not attend to a stimulus, the trial was repeated. If participants looked away in the middle of the videos, however, they were not repeated, as we aimed to test only responses to novel stimuli. Testing took place over 34 days for gorillas, 34 days for chimpanzees and 18 days for orangutans.

##### Humans

Adult participants were tested individually in a room with only the experimenter present. Participants first received written instructions to attentively watch the videos, and completed a demographic questionnaire on their linguistic background. Human adult participants also viewed all stimuli with ape agents first, followed by all stimuli with human agents. However, unlike for the ape participants, stimuli were presented in blocks (two blocks of 26 ape videos, followed by two blocks of 16 human videos). Short breaks were offered between blocks.

Human infants were tested individually in a laboratory of the research unit Developmental Psychology: Infancy and Childhood, with only an experimenter and a caregiver present. Before testing, parents completed a short questionnaire on the languages their child was exposed to, as an additional screening for any Ergative language experience. Infants were then seated on their parent’s lap in front of the eye tracker during testing, or in an infant carrier car seat, depending on which was more comfortable for them. To account for their decreased attention span, infants were presented with only half the stimuli seen by the ape and adult human participants (26 ape and 16 human videos; see Supplementary Material Table 1). To counter the reduction in number of trials compared with adults and apes, we increased the sample size accordingly. This included excluding any actions that might be considered unsuitable for infants, such as *hit, kick, pinch ear*.

Testing started with a short attention getter of dancing fruit followed by a 2-point calibration. Stimuli were presented in blocks (two blocks of 13 ape videos and two blocks of 8 human videos, presented in alternating order) following the calibration. Calibration was not always immediately successful, in which case to retain the infant’s attention, we continued with the presentation of the first block before attempting re-calibration. To ensure that this did not impact the overall number of human or ape stimuli viewed, the first block presented either ape or human videos first, in a counterbalanced order. If calibration was successful prior to the first block, at the start of the second block a 4-point validation was shown; otherwise, calibration was repeated. Each block started with a “practice” video, depicting either a nonhuman ape (ape blocks) or human (human block).

At the end of the session, adult participants or the infants’ caregivers were debriefed and given the chance to ask questions. For both adults and infants, sessions lasted approximately 30 minutes, including instructions and calibration.

#### Data processing

In order to determine what variables influence gaze, we coded the videos for perceptual and behavioral details. This included: 1) time at which the action of the agent started, 2) whether the agent or patient moved more, 3) whether tool use was involved, 4) whether food was involved, 5) whether at any point the agent or patient was camera facing, and 6) whether at any point the agent or patient had direct gaze with the camera. For details of the coding ethogram and interrater reliability, see Supplementary Material Table 2.

Areas of interest for agents and patients in the videos were manually defined in the Tobii Pro Lab software (see online Supplementary Material for examples). The eye-tracking data were then further preprocessed in R (58). Fixations were defined using the Tobii I-VT fixation filter algorithm. For each trial, we subsumed consecutive fixations to each area of interest into gazes, linearly interpolating gaps of up to 266 ms (80 eye tracker samples) between fixations to account for saccades and short periods of potential track-loss (57). Latency (in ms) of the first gaze to either the agent or the patient was extracted. Gazes were then aggregated into time bins to reduce temporal auto-correlation (59,60). Bins were defined relative to the length of each stimulus video, so that each time bin encompassed the fixations from a segment spanning 5% of the stimulus length. This binning normalizes stimulus length and thus makes videos of different durations comparable.

Trials were excluded from analyses if the first gaze into a stimulus occurred later than 1000 ms after video onset or if no fixations were detected. Trials from apes were removed if they looked at the area of the video before it started playing on screen (as observed by the experimenter). For the infants that were not fully excluded, trials were excluded due to poor or missing calibrations or because the infants were not attending to the stimuli. Trials in which participants never fixated on the agent or patient or in which none or less than 50 ms of eye tracker samples were registered before the first fixation was detected were also excluded. In addition, three video stimuli presented to human adults, and two video stimuli presented to apes and human adults, were excluded from analyses due to technical errors with the videos; one video stimulus presented to apes and human adults was also excluded, because the agent and patient were considered too ambiguous. Overall, 3537 (out of 5628 total trials) trials were included in the statistical analyses.

#### Statistical analyses

The data were analyzed in R (version 4.2.1) with Bayesian multilevel models on the single trial level, using the RStan package (58,61), which fits Bayesian models using a Hamiltonian Markov chain Monte Carlo algorithm. Markov chain convergence was assessed using standard diagnostics (number of effective samples, the Gelman-Rubin diagnostic, and visual inspection of trace plots).

Our outcome variable was the AOI (e.g., Agent, Patient, or Other) that a participant was gazing at during time 𝑡, where 𝑡 ∈ [*1*,*20*] indexes the 5% interval of the total duration of a trial such that 𝑡 = *1* corresponds to the time interval [*0*,*0*.*05*), 𝑡 = *2* corresponds to [*0*.*05*,*0*.*10*), etc. In principle, a participant could gaze at multiple AOIs during an interval. However, given that these intervals correspond to approximately 300 ms (mean = 319.939, sd = 103.783; varies by stimulus), and the average duration of eye fixations in humans is 200-300 ms (62), a time bin should usually capture only 1 unique fixation event. As such, heterogeneity within such a short time interval may be best understood as measurement error.

That said, the choice of a 5% window is somewhat arbitrary. Ideally, we would choose a binning window length that is short enough not to average over true fixations, while not being so short that adjacent samples are perfectly auto-correlated, a ubiquitous problem in eye-tracking research and time series modeling more generally. We can quantify the information gained by decreasing interval length by looking at the correlation between the data at 5% intervals vs 2.5% intervals–a doubling of temporal resolution. We found that, averaging over the three AOIs, the correlation between the 2.5% and 5% binned variables was 𝜌 = *0*.*929*. This suggested that little information would be gained by further increases in temporal resolution, and thus we used 5% intervals to balance data loss with computational burden.

Another source of heterogeneity within time intervals (i.e., a mixture of gazes between different AOIs) is that, for some videos, the Agent and Patient AOIs overlap on the screen, such that it is possible for a participant to be gazing at both simultaneously. Thus, even if there were no measurement error we would sometimes observe a mixture of AOI gazes during a short time window. To propagate uncertainty in our model due to both measurement error and overlapping AOIs on the screen, we weight the probability of gazing at each AOI (a latent state, denoted by 𝐾, where 𝑘 = *1* indicates Agent, 𝑘 = *2* indicates Patient, and 𝑘 = *3* indicates Other) by the observed proportion of gazes for each AOI during an interval 𝑡, denoted 𝑌 where 𝑦_𝑡,𝑘=_ *_1_* indicates the proportion of agent gazes during 𝑡, 𝑦_[𝑡,𝑘=_ *_2_*_]_ indicates the proportion of patient gazes, and 𝑦_[𝑡,𝑘=_ *_3_*_]_ indicates the proportion of other gazes.

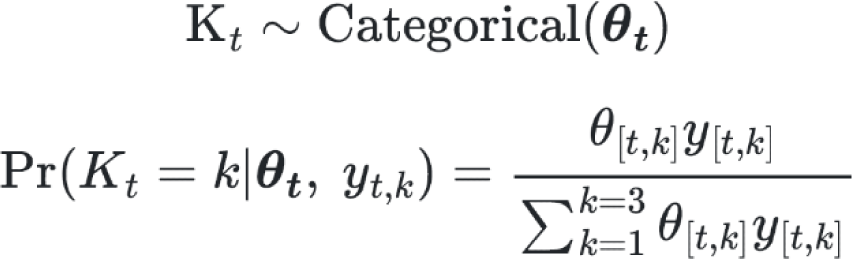

Where 𝜃_𝑡_ is a vector of probabilities for each latent AOI state, (e.g., 𝜃_[𝑡,𝑘=_ *_1_*_]_ is the model probability of Agent during time 𝑡) such that 𝜃_𝑡_ sums to 1. With this approach, we were able to model AOI gaze as a latent categorical variable while propagating the uncertainty that is inherent to the data generating process (i.e., measurement error).

We can then make 𝜃_𝑡_ a function of predictors in a generalized linear model using the softmax link function. As such, there were 𝑚𝑎𝑥(𝐾) − *1* linear models for the latent “scores”, denoted 𝑠, which the softmax function maps to the vector of probabilities 𝜃_𝑡_. “Other” was set as the reference category, so that 𝑠_[𝑡,𝑘=*3*]_ = *0* for all 𝑡.

To adjust for potential confounding factors, the model included as predictors: the ratio between the mean agent and the mean patient AOI size between the video stimulus start and the onset of the first gaze (Supplementary Material Figure 5); the category of the depicted event (“social” events with both an ape or human agent and patient or events where an ape or human agent interacted with an inanimate object, or with food); difference in movement between agent and patient (coding which moved more; Supplementary Material Figure 6); the participants’ species and the depicted species, as well as group-level predictors for the individual stimuli and participants. In most stimuli, the action of interest started immediately at the onset of video presentation. In 25 stimuli, however, the action started only after video onset (on average 0.24 seconds after); for these stimuli, the time points before the action started were excluded from the models.

Below we describe the linear model, first as a simplified version using natural language, and then a formal definition:

**Natural language definition**

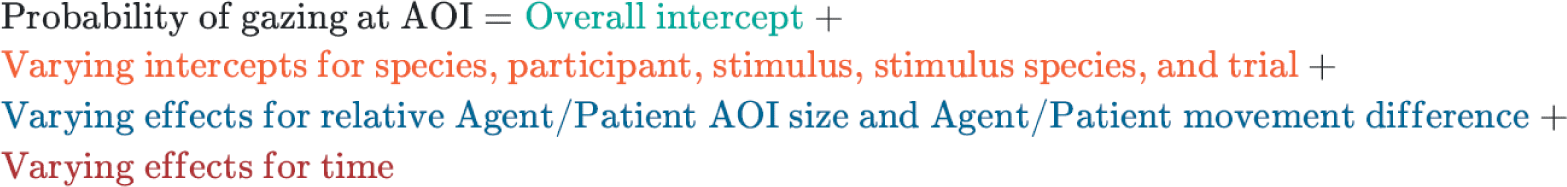

**Formal definition**

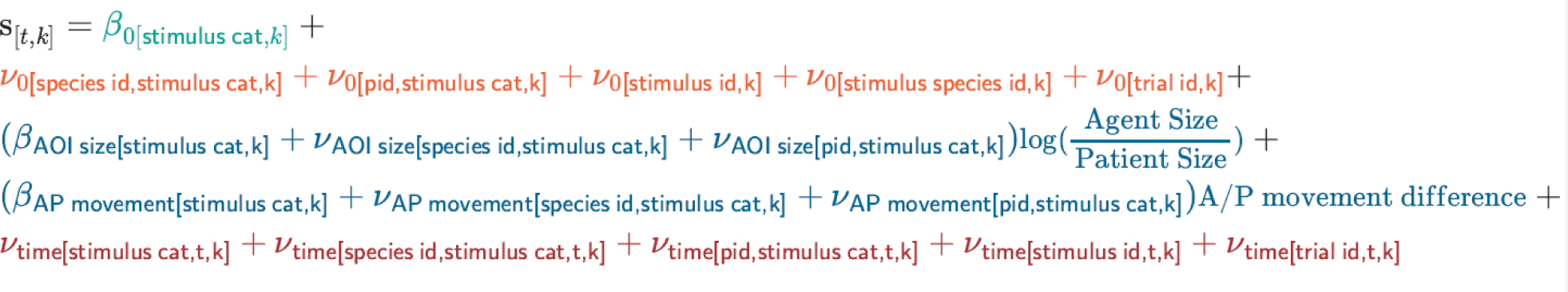

Where “stimulus cat” is an index of whether the video (stimulus) being viewed is inanimate (food), inanimate (non-food), or social. Stimulus ID indexes the identity of the specific video being viewed in a trial, while Stimulus species ID indexes the identity of the species featured in a video. 𝛽 indicates an overall (also known as “fixed”) effect, while 𝜈 indicates a varying (also known as “random” or “mixed”) effect. The varying effects for time co-vary according to a Gaussian process with an exponential kernel:

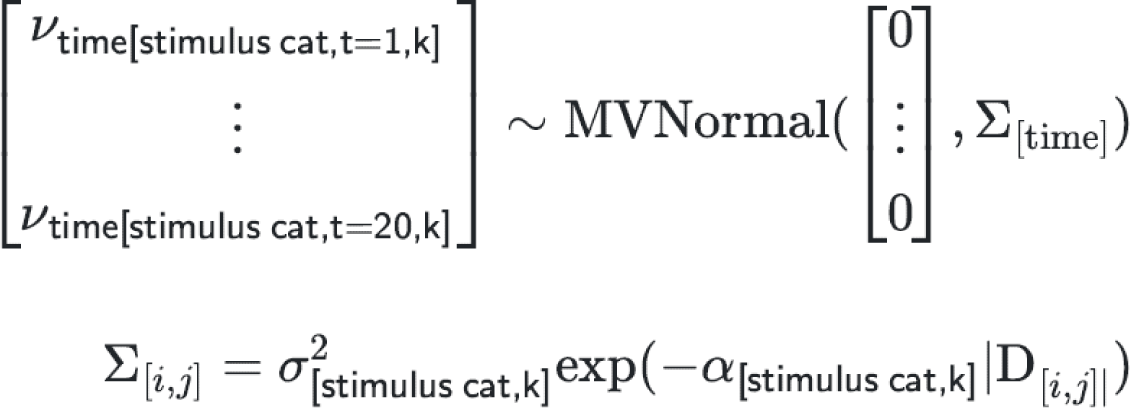

Where 𝐷 is a matrix of pairwise distances between time intervals. Adjacent intervals have a distance of 0.05 (corresponding to 5% time bins), intervals two steps apart have a distance of 0.1, and so on. The relative “wiggliness” or rigidity of these time functions are controlled by the parameter 𝛼 (large values of 𝛼 imply relatively wiggly functions that change rapidly, small values imply relatively rigid functions that change slowly). The magnitude of the functions (i.e., the extent to which they affect the latent scores) is controlled by 𝜎*^2^*.

The time varying effects for species, participant (pid), stimulus, and trial are modeled using hierarchical Gaussian processes, which means rather than estimate each time function for each group independently, the model estimates shared 𝛼 and 𝜎*^2^* parameters for each level. Thus, each species, participant, stimulus, and trial can have a unique time smooth, but these functions are regularized via partial pooling to reduce over fitting. As an additional layer of regularization given the high number of parameters in our model, we employed a Dirichlet decomposition for each set of varying effects (63,64). This means that we parameterized the variance of each vector of varying effects 𝜈 in terms of the total variance for that set (e.g., the total variance of the intercepts, or time effects) and a simplex 𝜙 consisting of level-specific variance components. For example, if 𝜎*^2^* = 2, and 𝜙_𝑝𝑖𝑑_ = *0*.*3*, then the variance for the participant-level effects is 2(0.3) = 0.6. By avoiding separate variance parameters for each level (i.e., separate 𝜎*^2^* priors for species, participants, stimuli, trials), we constrain the prior predictive variance.

We set generic, weakly-regularizing priors on all model parameters:

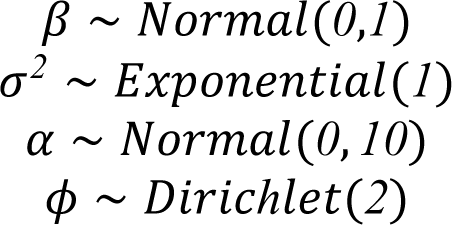

See Stan code for additional details on the implementation of this model.

### Trial Averaged Model Comparison

In addition to the time series model described above, we also fit two models designed to test whether there was an overall difference in the frequency of Agents vs Patients gazes, averaging within trials. To do this, we calculated the trial-level proportion of gazes to each AOI, shifting our response variable from a Categorical distribution (the AOI category at each time point) to a Dirichlet distribution (the proportion of gazes directed to Agent, Patient, and Other over the entire trial, where the three components sum to 1). For these models we retained the same multilevel model structure and covariates as the time series analysis, omitting only the time-varying components and the trial-level random intercepts (because the unit of analysis is now the trial).

Where y is a vector containing the trial level proportions of gazes at each AOI, α is the parameter vector of the Dirichlet distribution, which we re-parameterize in terms of mean vector μ and dispersion parameter φ. We specify linear models for each element in η, which represent the latent-scale expected proportion of gazes for each AOI.

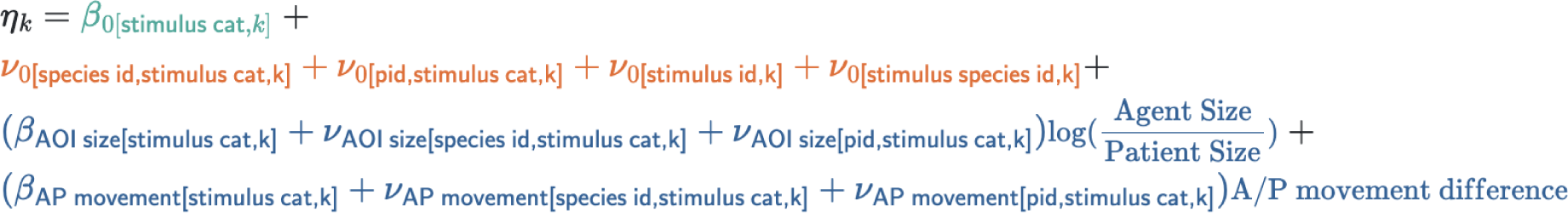

As before, we set weakly-regularizing priors on all model parameters:

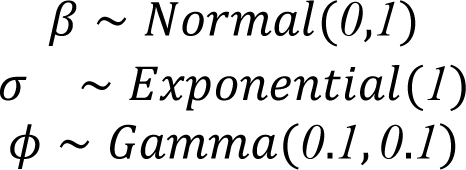

To set up our null model (equal gazes to Agents and Patients, conditional on covariates), we used the same model structure as just described, except that we forced the linear model for the Agent AOI to be equal to the Patient AOI rather than estimate each of them. This forces the two AOIs to always have the same conditional probability, which is what we would expect if Agents and Patients were exchangeable.

After fitting both the null and alternative models, we compared them using approximate leave-one-out cross-validation via the expected pointwise predictive density (ELPD). The ELPD is approximated by Pareto-smoothed importance sampling (PSIS) of the posterior (65). To assess relative model performance, we used model stacking, which allocates weights to models such that they jointly maximize predictive accuracy (66). These stacking weights are interpreted as the probability that a model will perform better than any other model in the comparison set (in this case, just the null and alternative models). ELPD and stacking weights were computed using the ’loo’ R package (67).

## Supplementary text

### Ape testing

#### Habituation procedure

Initial habituation took part in two phases. In the first phase, apes were shown the liquid reward to encourage them to approach and inspect the screen. If they attended to the screen then a video would be played. Stimuli were presented using either Tobii Studio (for gorillas) or Tobii Pro Lab (for chimpanzees and orangutans). In the first phase, 75 videos depicting varied species, actions and interactions were played, to engage participants, and to determine interest and emotional response to different types of videos. The goal was to ensure that participants were comfortable to approach the screen and attend to the videos, and that they kept a good position during viewing. After successfully viewing all footage of the first phase, they were then shown footage from phase 2. This consisted of 16 videos depicting a variety of species, selected for their causal structure. We used gaze responses to this footage to help determine AOI regions for the test footage, and to finalize the most appropriate footage for testing. Habituation and calibration took place over two months for the gorillas, three months for the chimpanzees, and four months for the orangutans.

### Test footage collation

Footage of apes were collated from a combination of captive and wild footage from our own archives, other researchers and the Disney movie Chimpanzee, as well as filmed for the purposes of this study at Zurich Zoo (see Supplementary Material Table 1 for full list). Natural scenes depicted footage of dyadic social interactions such as play, grooming, approaching and infant suckling, as well as inanimate interactions such as foraging, eating, using enrichment and tool use. We chose stimuli of relatively neutral content so as to avoid arousal responses which could distract participants.

**Supplementary Material Figure 1.**
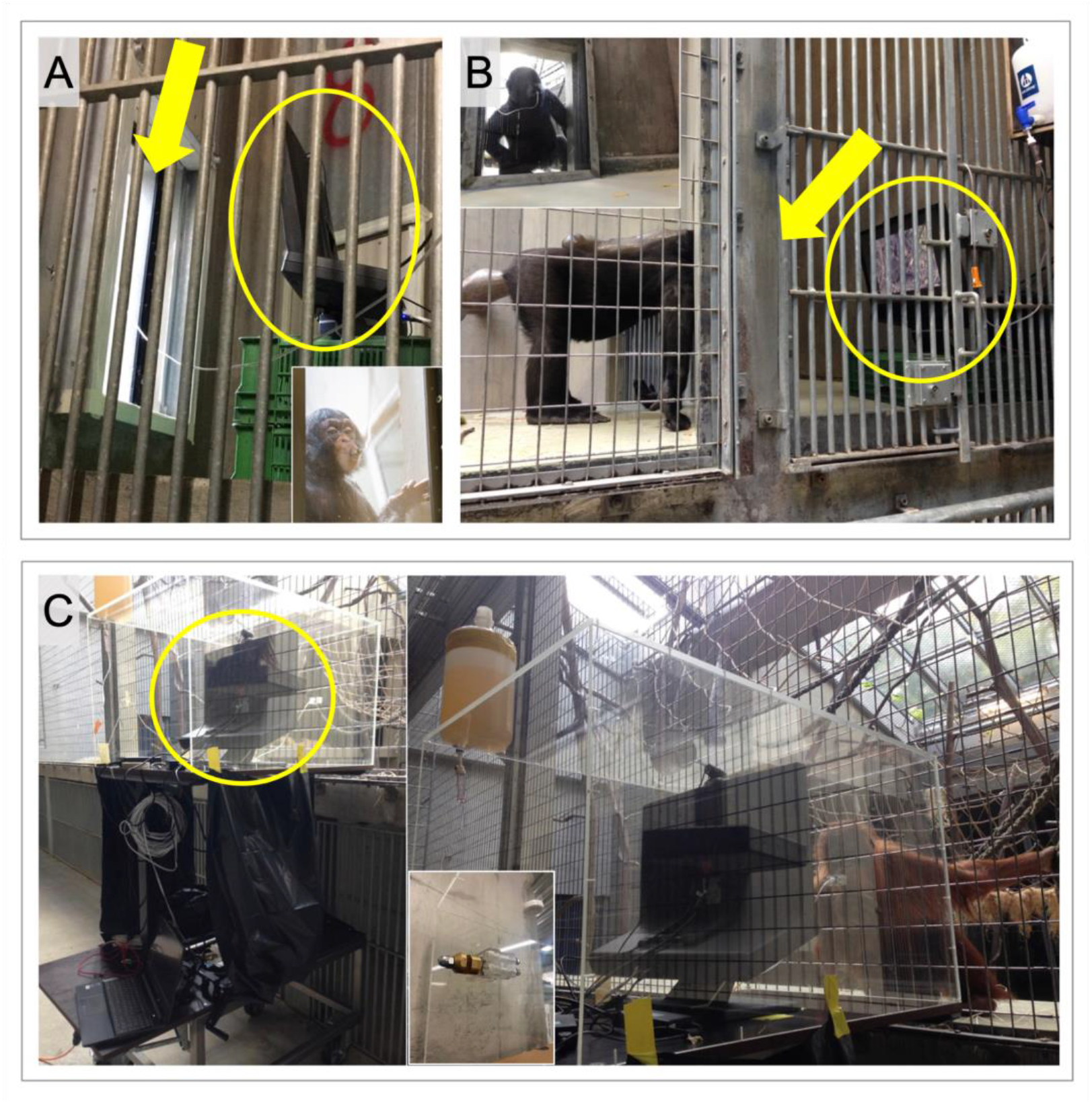
Eye-tracking setups at Basel Zoo. A: fixed setup for chimpanzees. B fixed setup for gorillas. C: mobile setup for orangutans; indent shows drinking nozzle with hook to attach to mesh. Circles indicate display monitor and arrows indicate viewing window.

### Variable coding scheme

The coding scheme is detailed in Supplementary Material Table 2. To determine interrater reliability for these variables, coding was conducted by two coders. For the variables time action starts and agent-patient movement difference, both coders coded and discussed differences for 20 video clips; they then independently coded an additional 21 test clips (25%). For the variables direct gaze at camera and orientation towards camera, due to the low number of instances of each behavior, both coders coded all videos. Interrater reliability was calculated using Cohen’s kappa. Values are reported next to each variable in the table.

**Supplementary Material Table 1.**
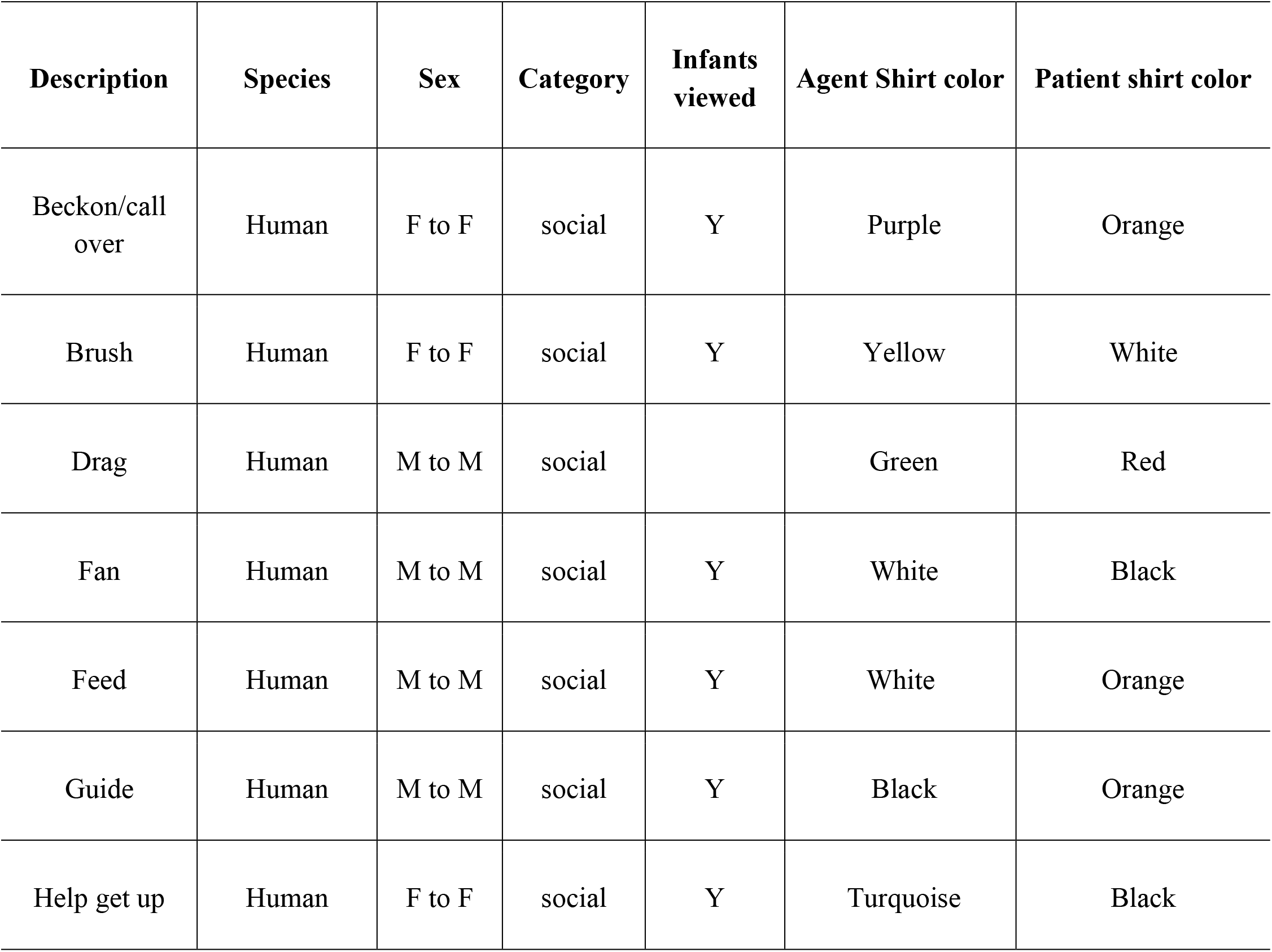
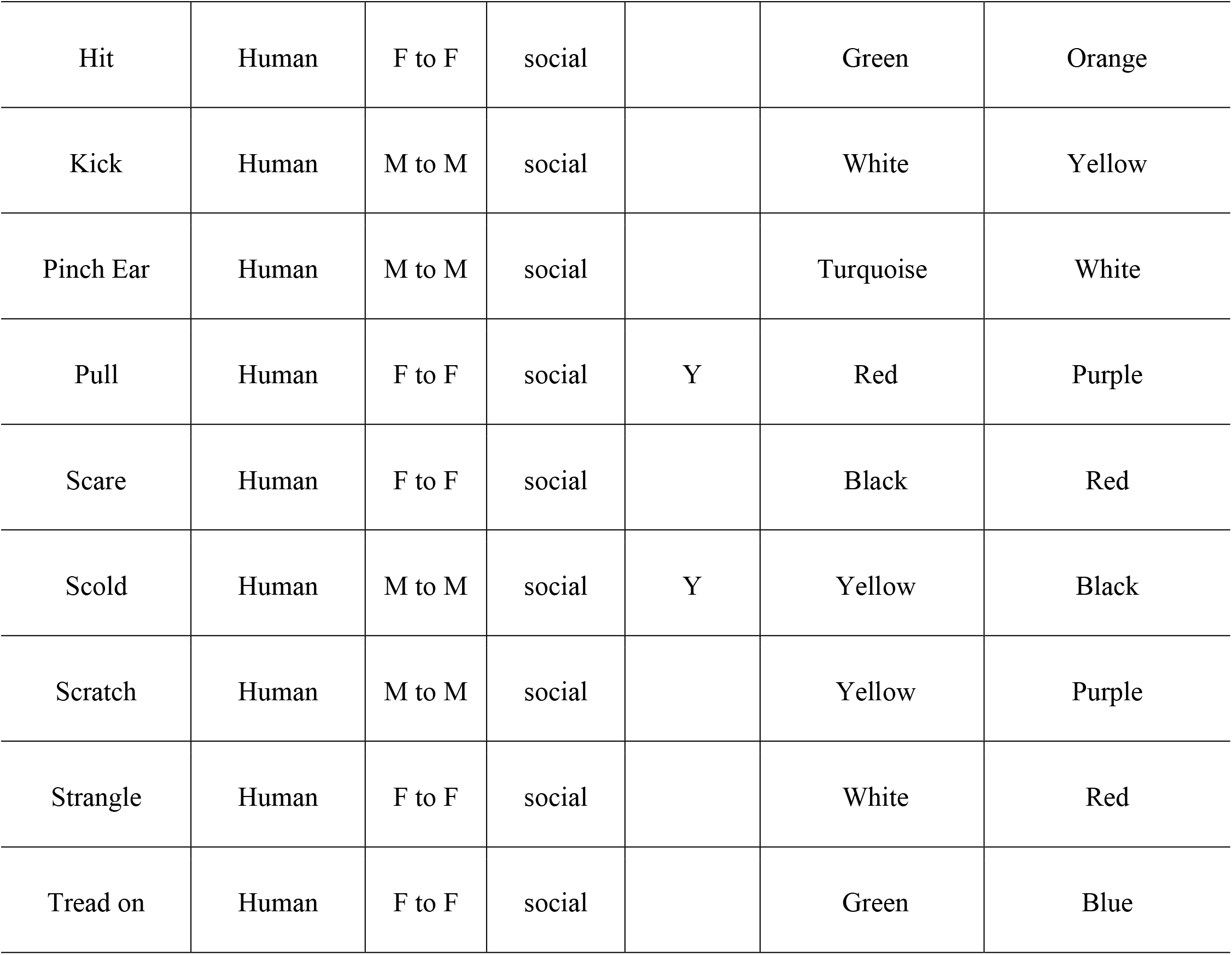
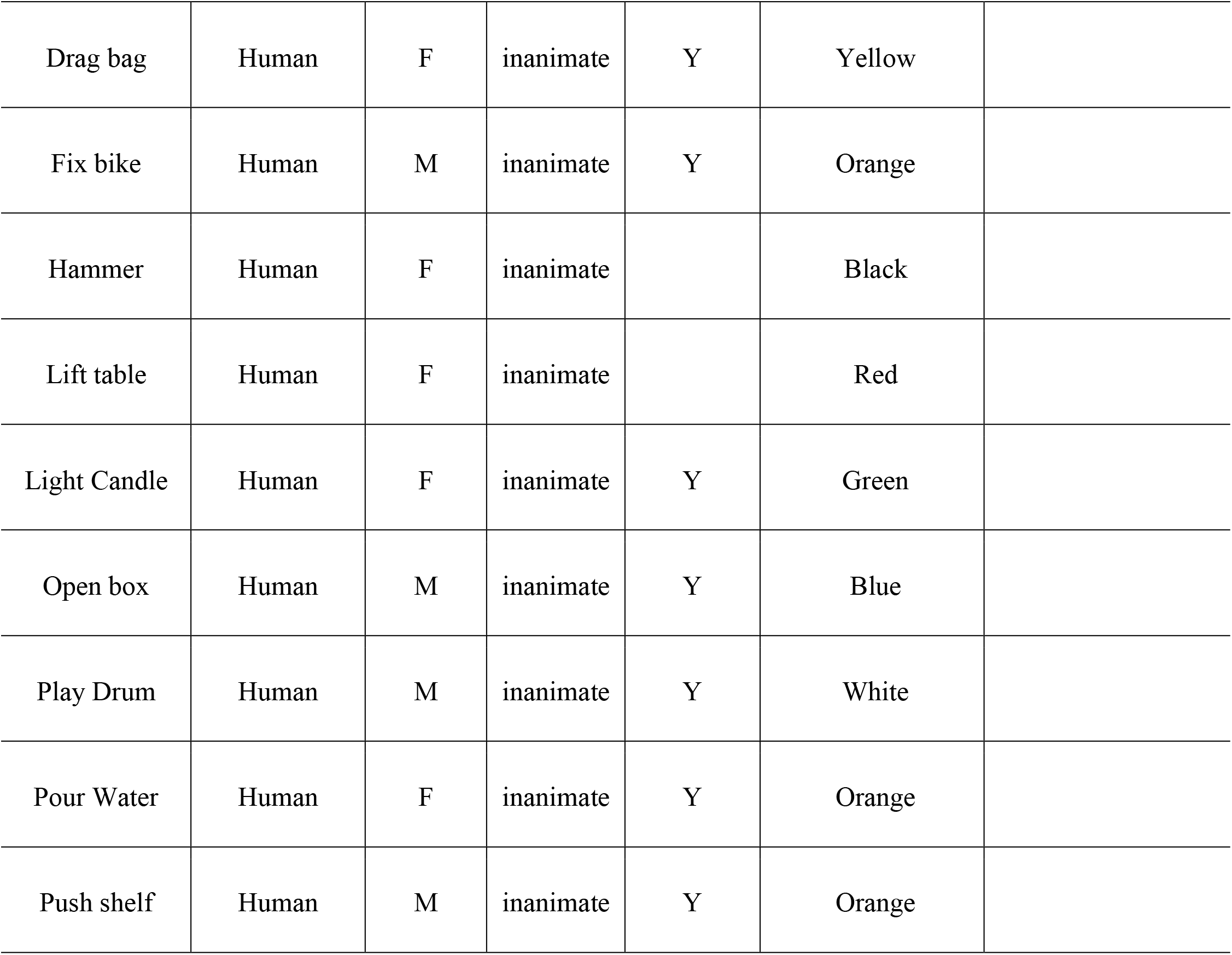
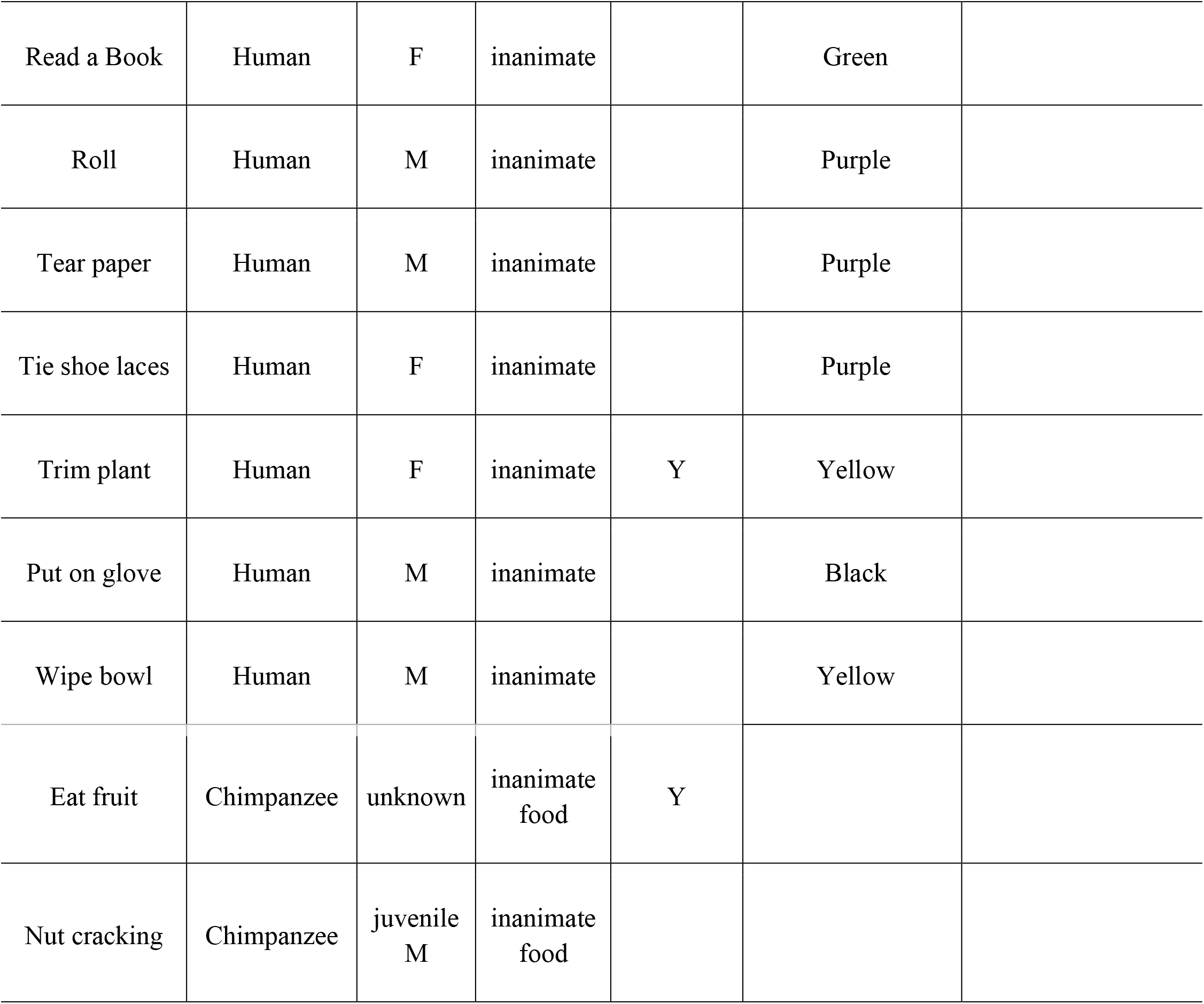
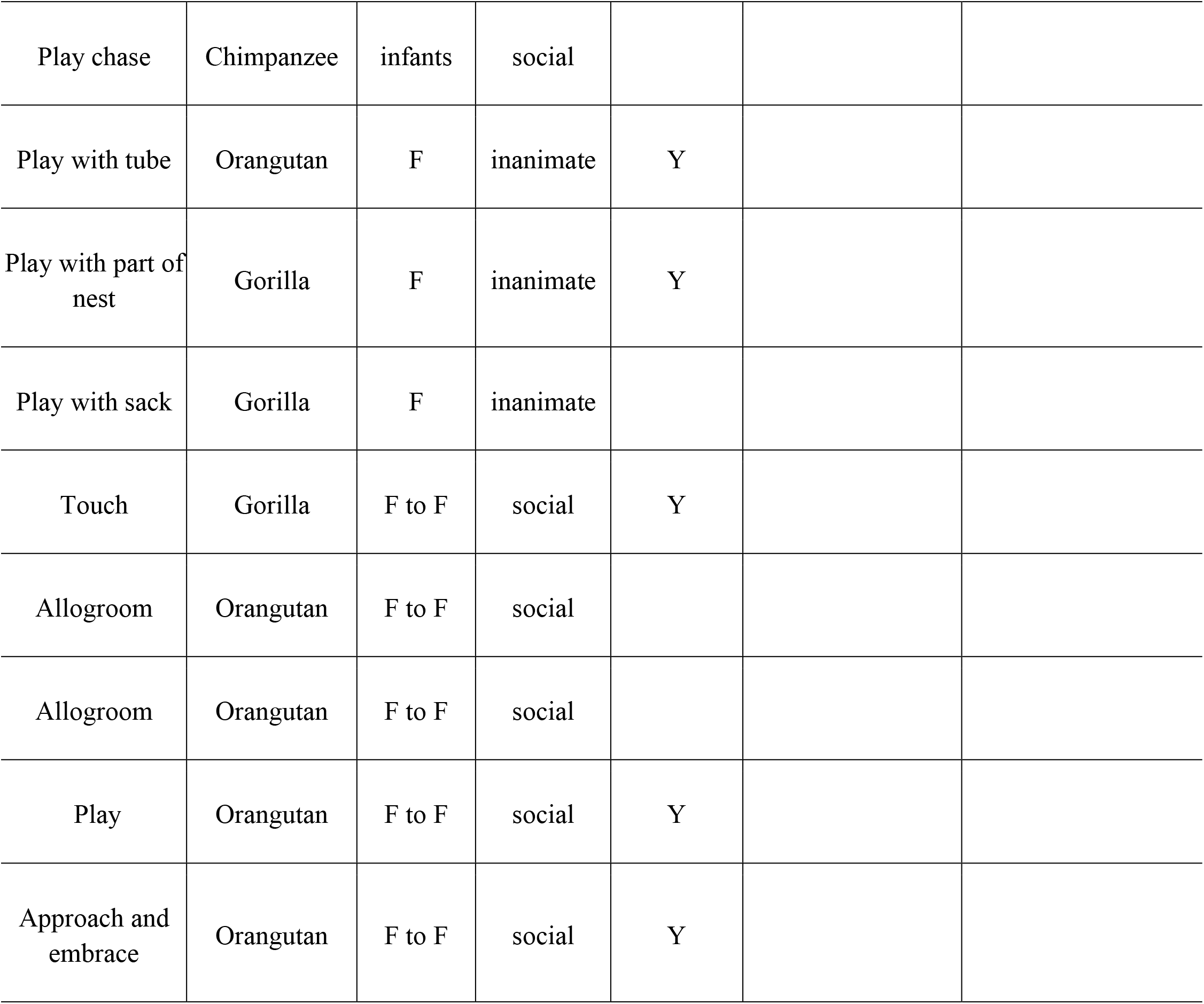
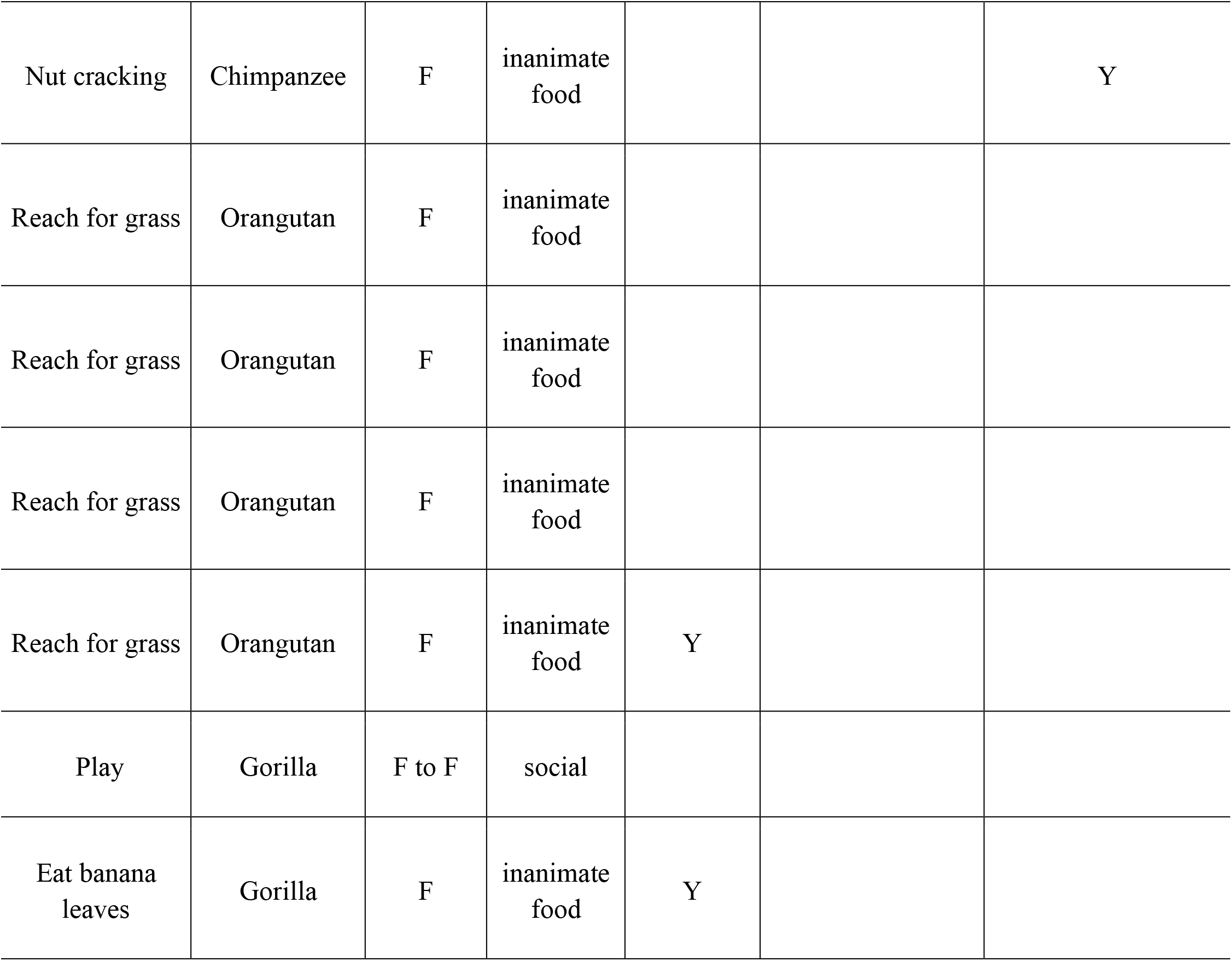
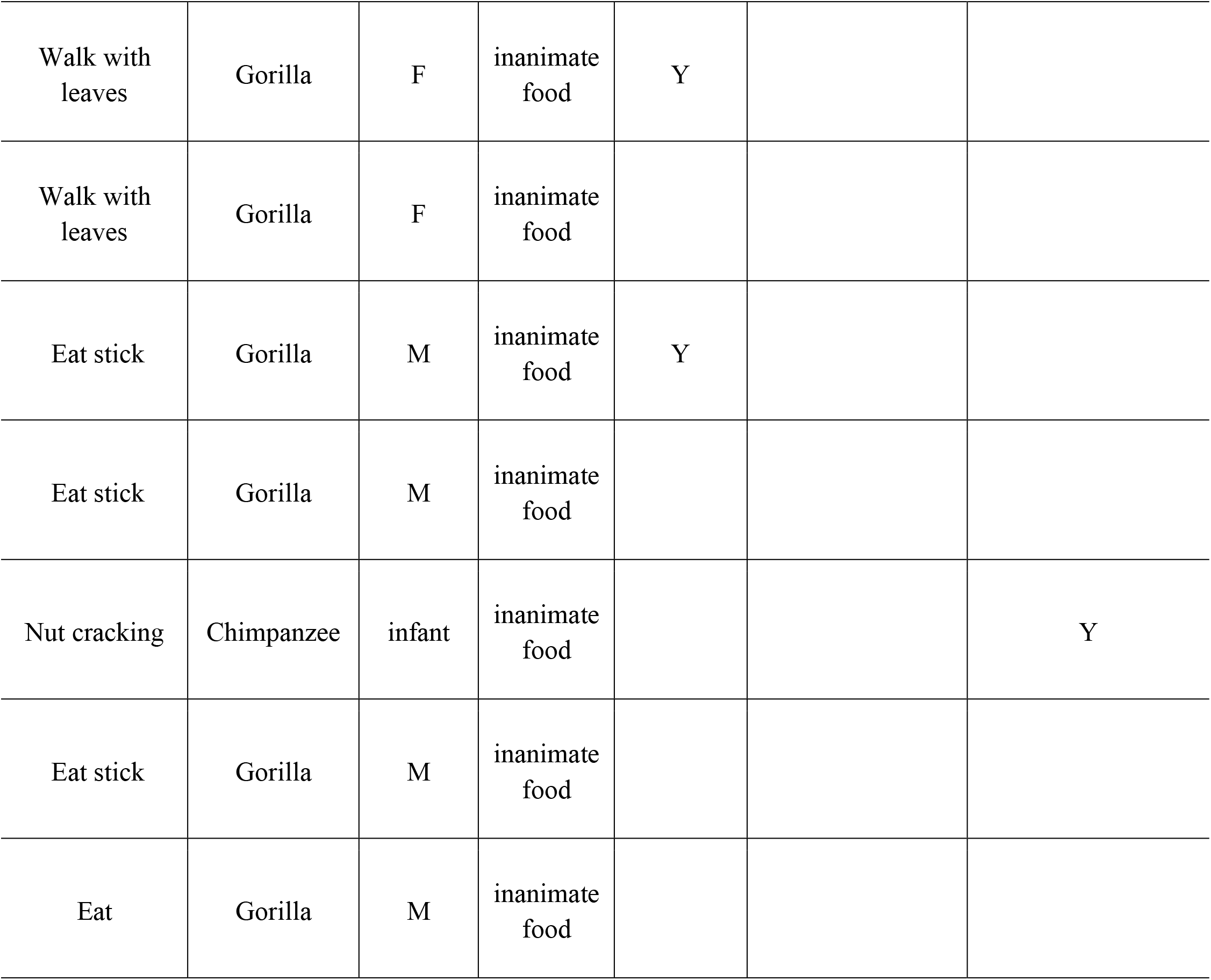
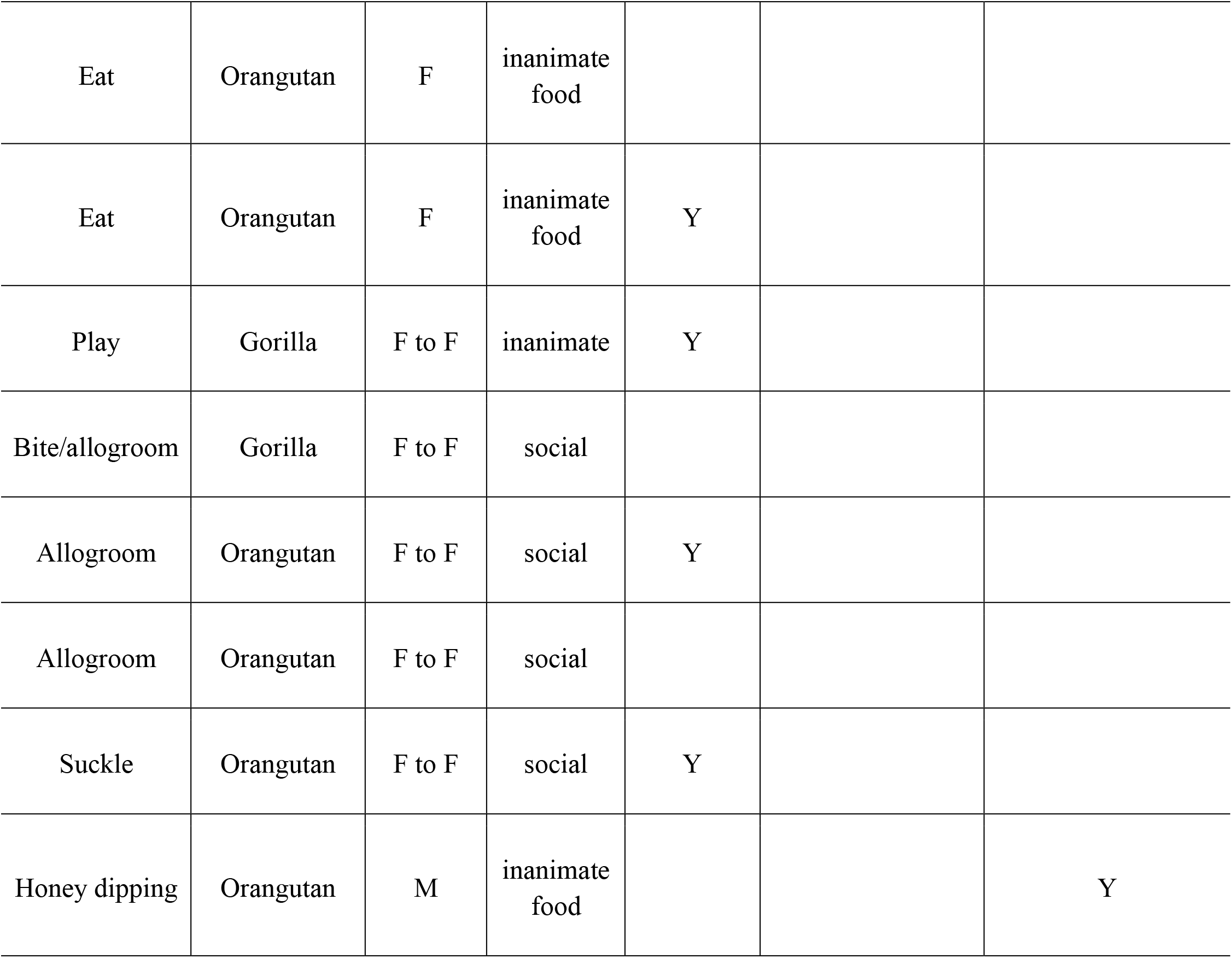
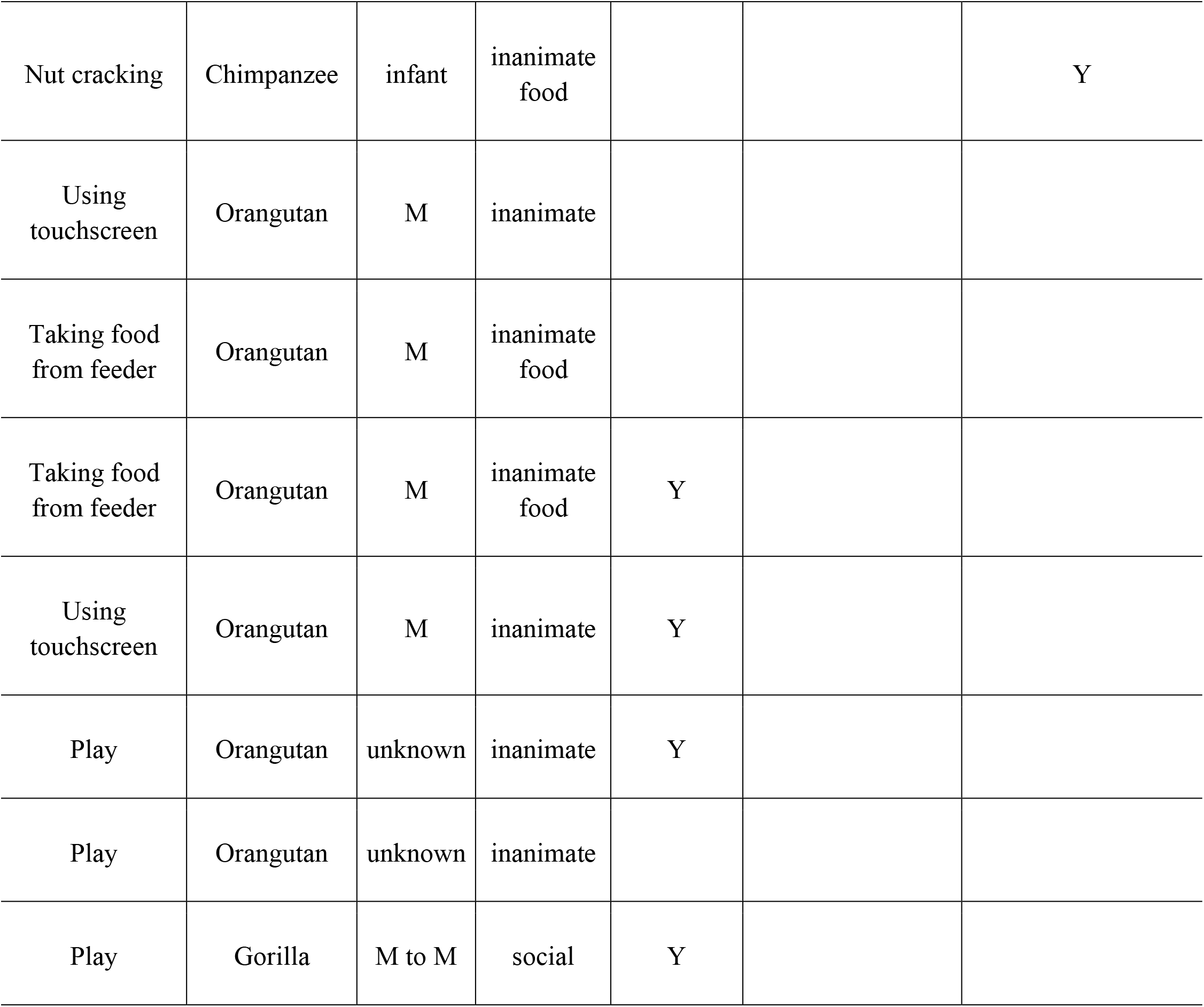
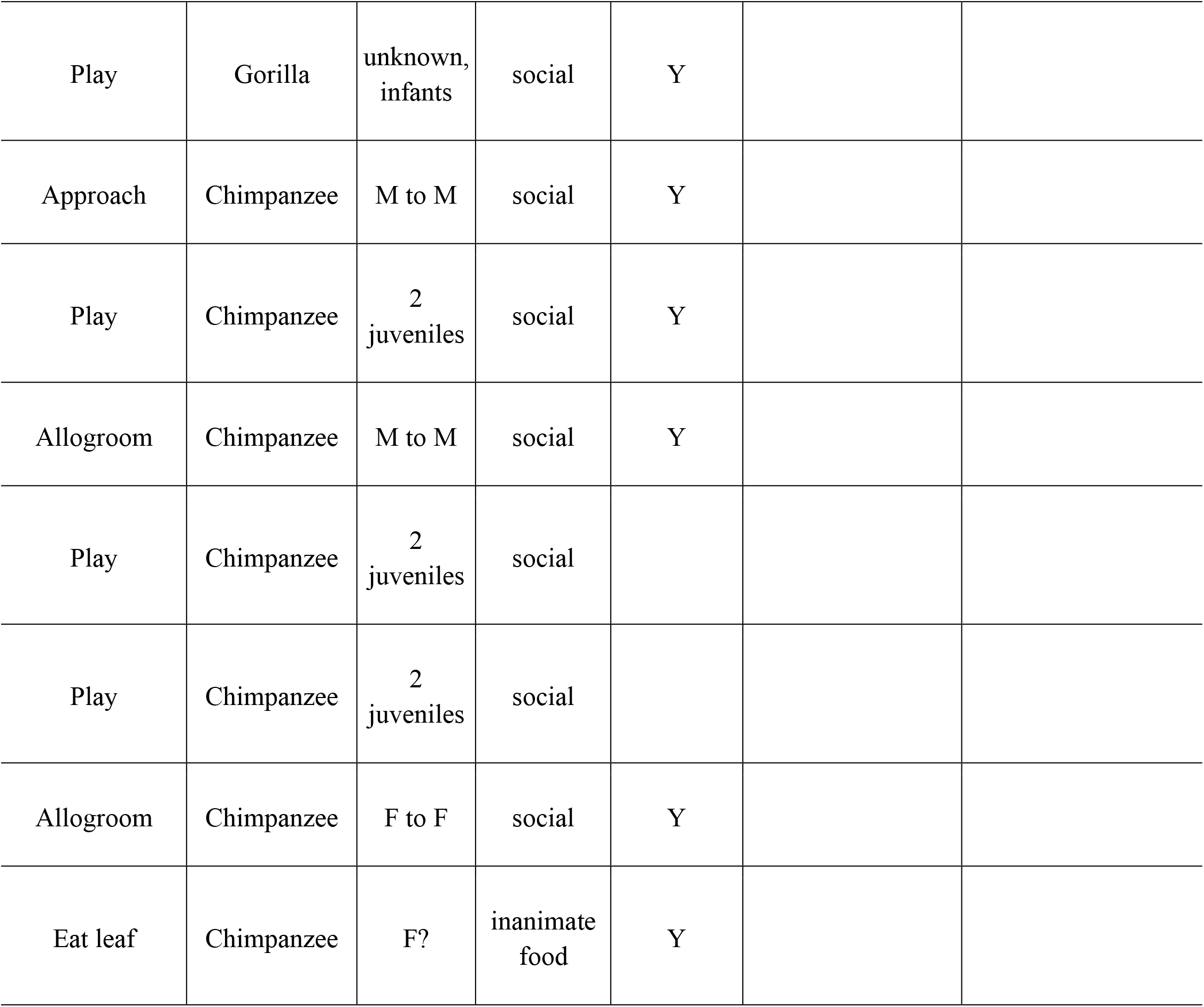
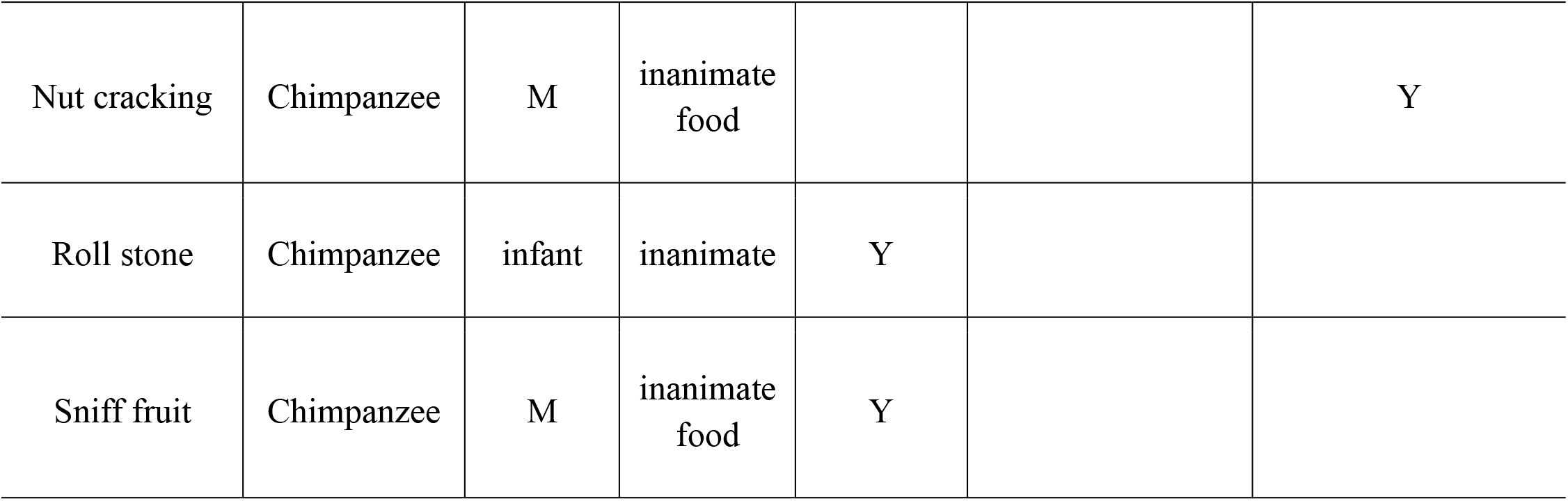
Full list of video footage presented

**Supplementary Material Table 2.**
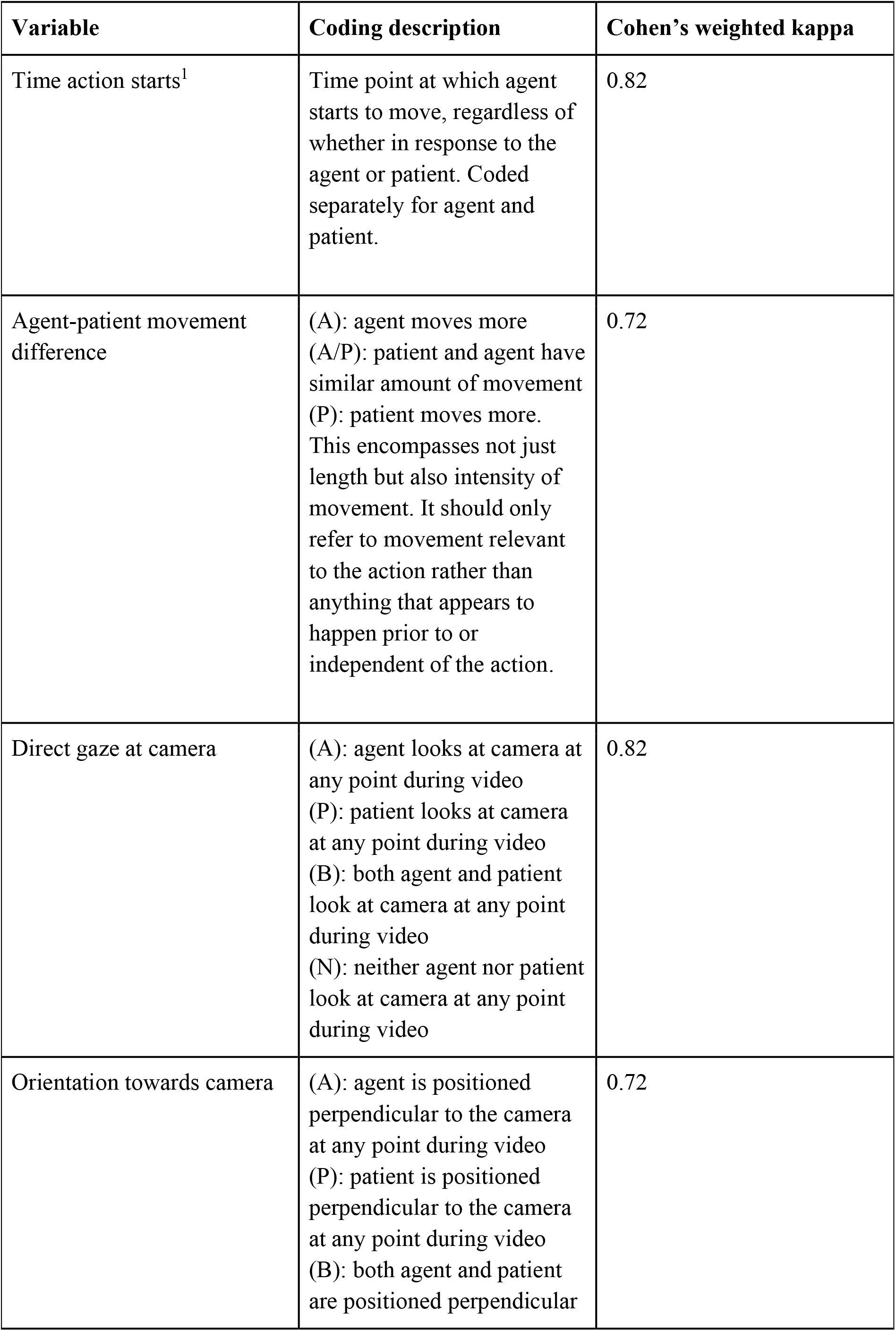
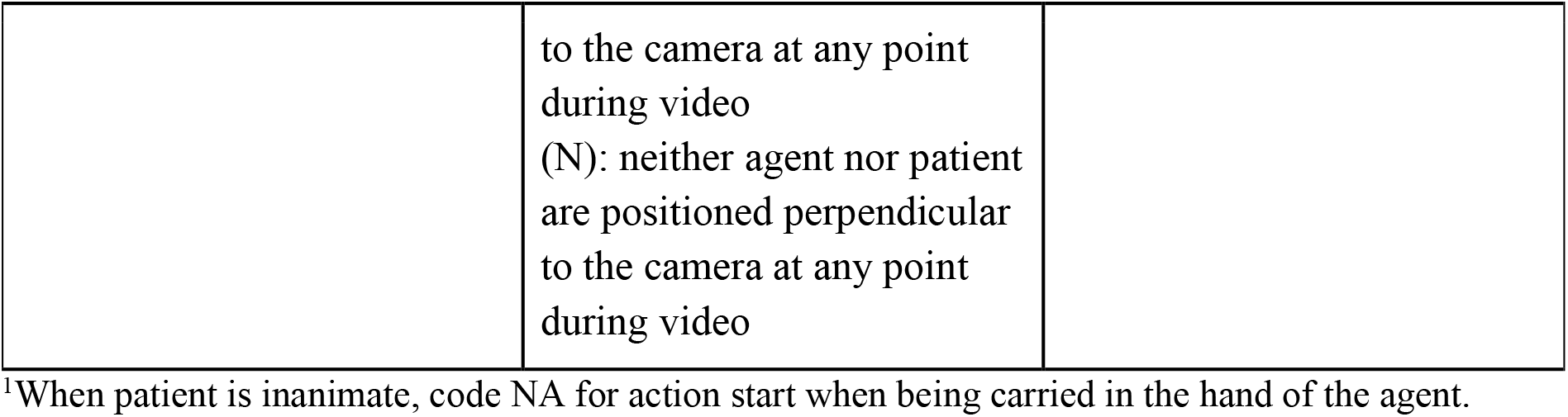
Variables coded from video stimuli

## Analyses and Results

**Supplementary Material Figure 2:**
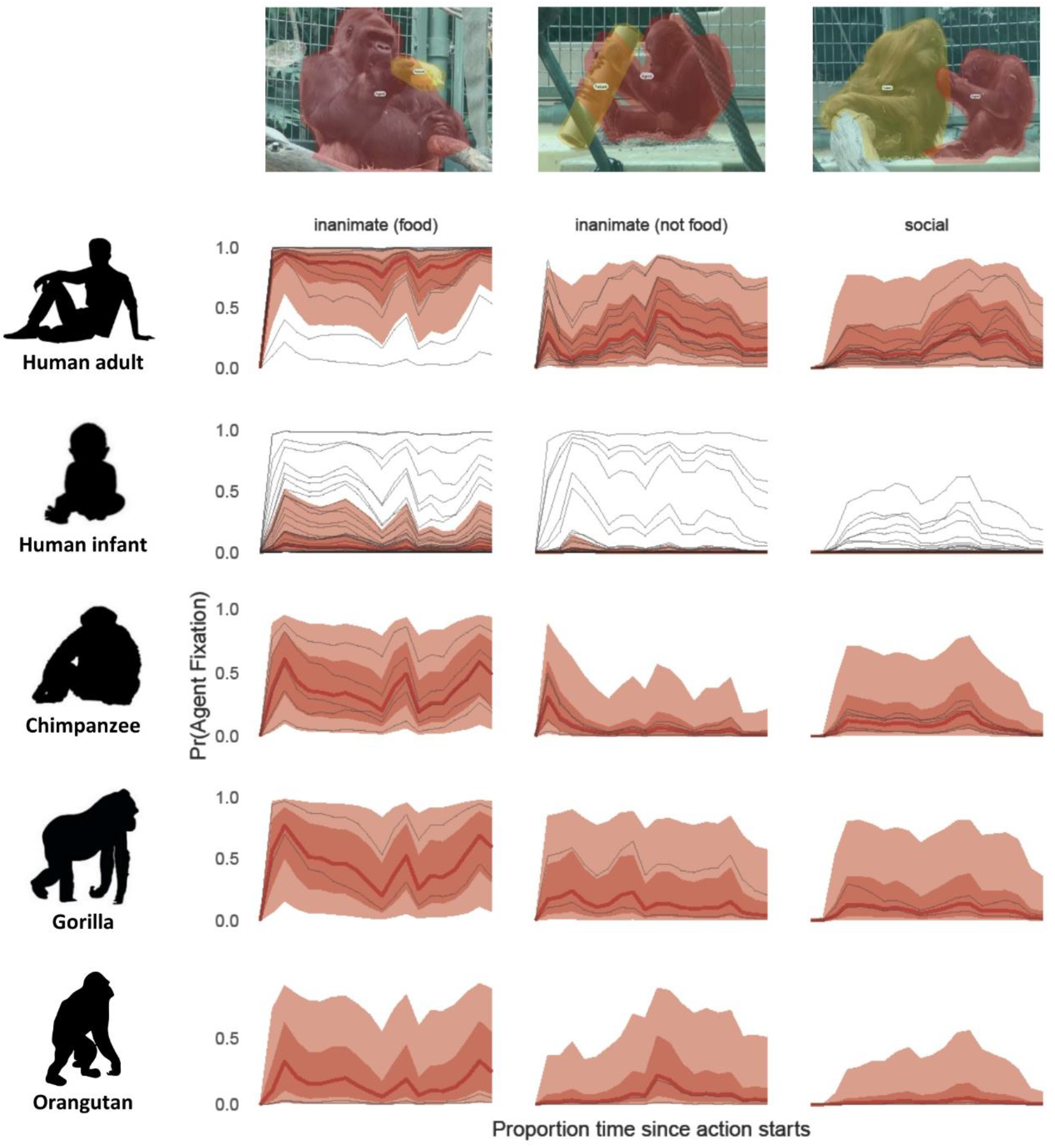
Time course of gazes towards agent in the videos, predicted with a categorical Bayesian multilevel model. Thick lines represent the grand mean, thin lines represent individual participants. Time point 0 on the x-axis indicates action start time, normalized across stimuli; fixations were aggregated into 5% time bins; shaded ribbons indicate 50 and 90% credible intervals, respectively.

**Supplementary Material Figure 3:**
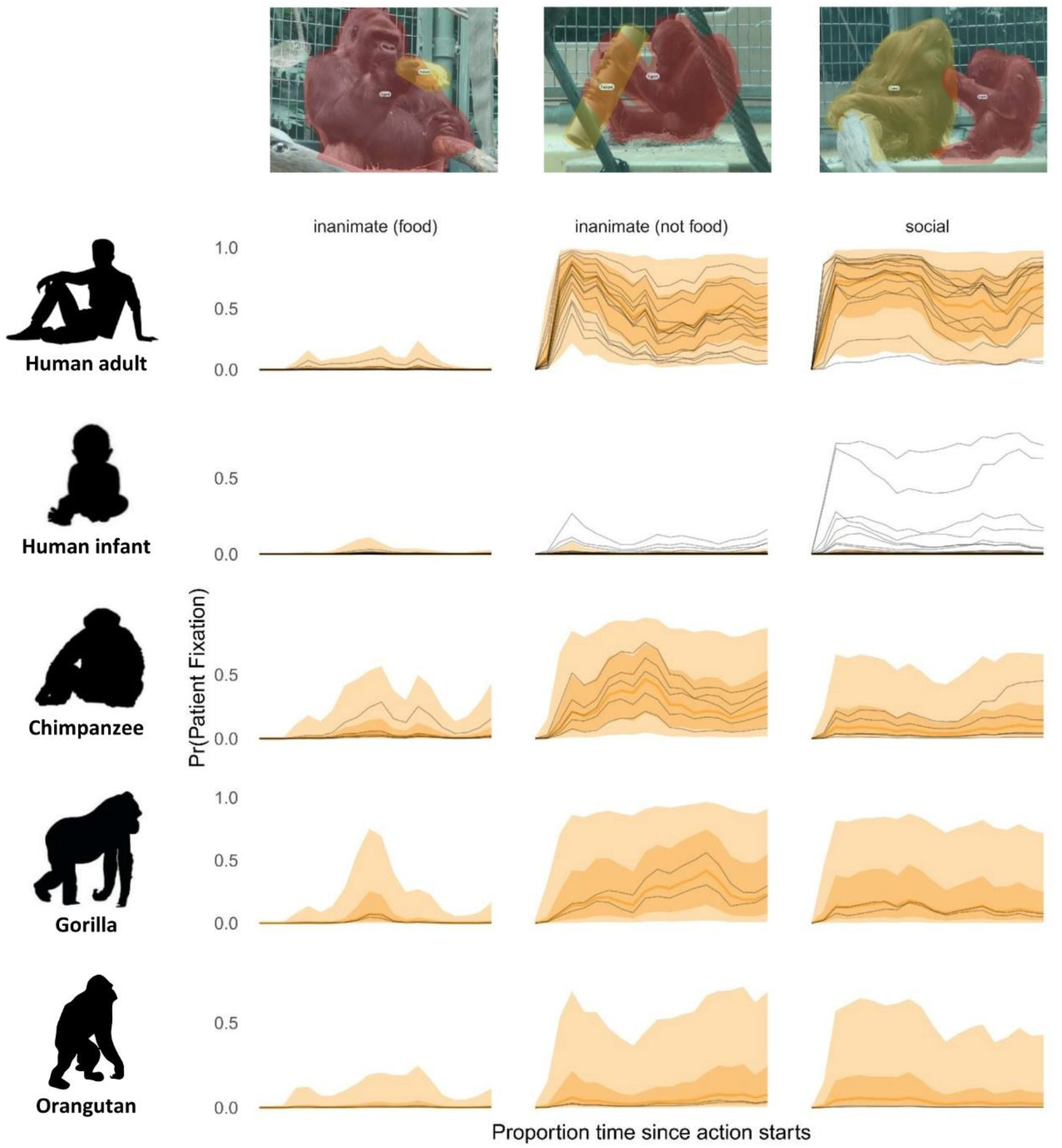
Time course of gazes towards patient in the videos, predicted with a categorical Bayesian multilevel model. Thick lines represent the grand mean, thin lines represent individual participants. Time point 0 on the x-axis indicates action start time, normalized across stimuli; fixations were aggregated into 5% time bins; shaded ribbons indicate 50 and 90% credible intervals, respectively.

**Supplementary Material Figure 4:**
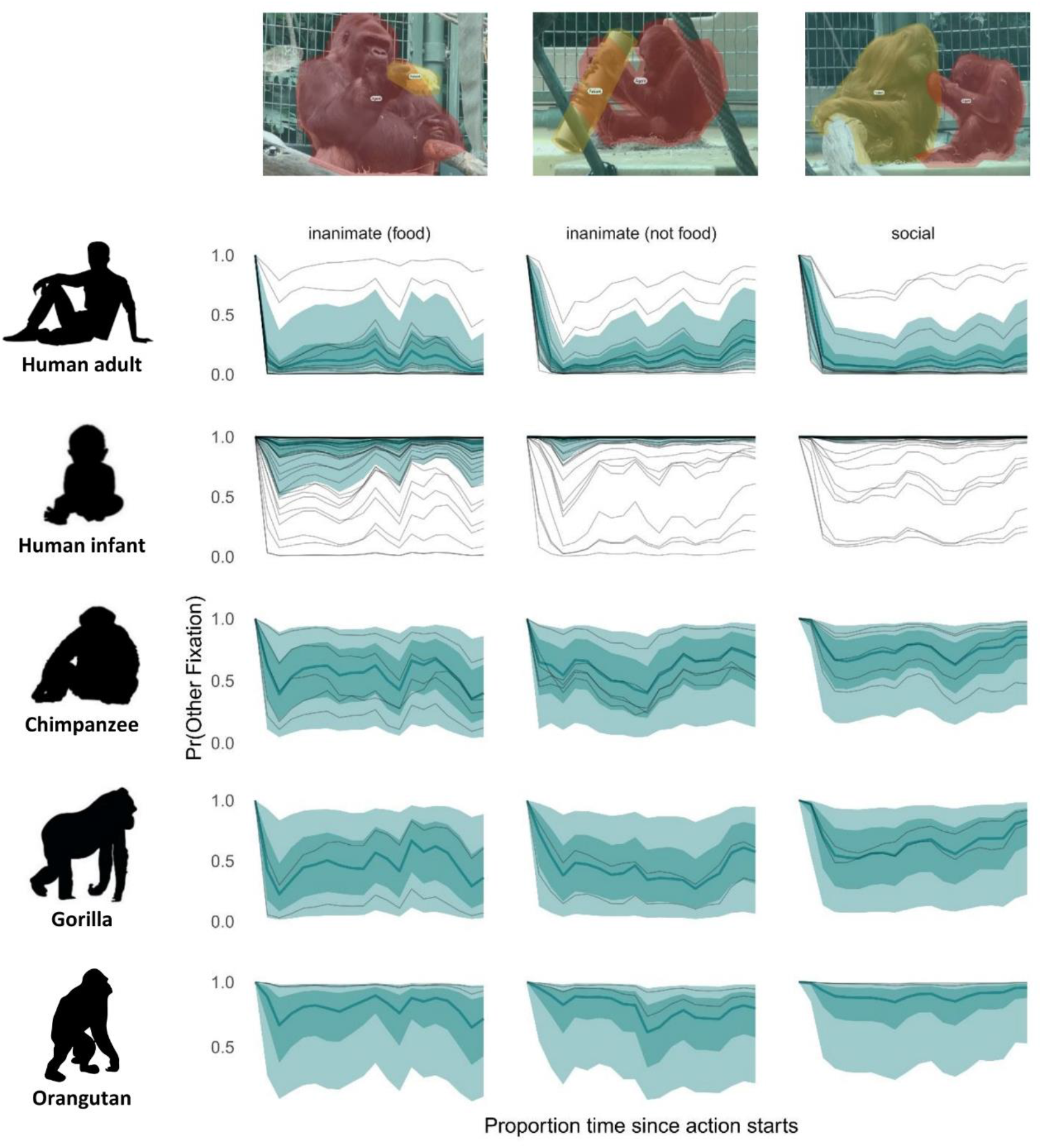
Time course of gazes towards other information in the videos, predicted with a categorical Bayesian multilevel model. Thick lines represent the grand mean, thin lines represent individual participants. Time point 0 on the x-axis indicates action start time, normalized across stimuli; fixations were aggregated into 5% time bins; shaded ribbons indicate 50 and 90% credible intervals, respectively.

**Supplementary Material Figure 5:**
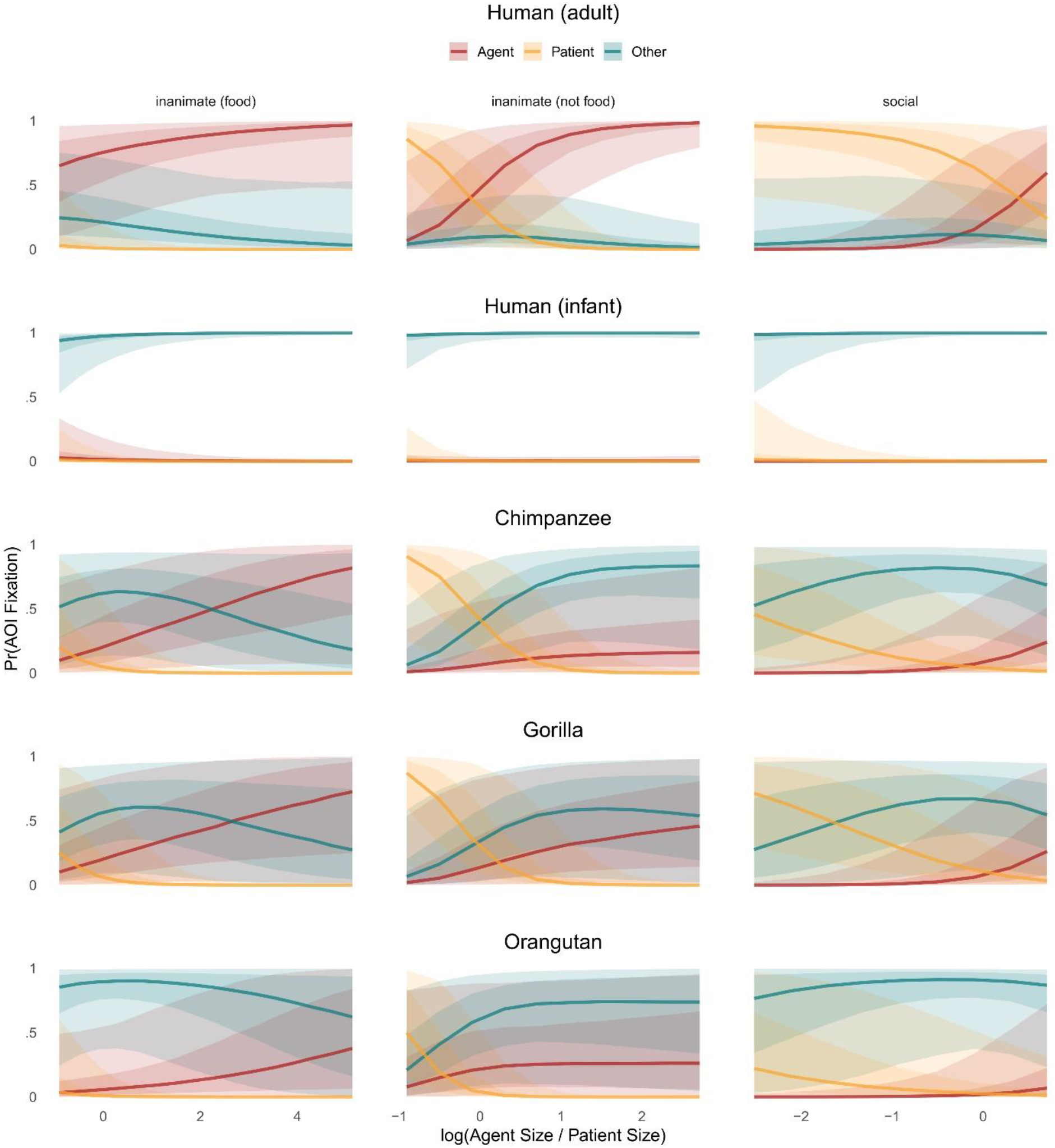
AOI fixation probability as a function of AOI size. AOIs are depicted by color: agent (red), patient (orange), and other information (turquoise).

**Supplementary Material Figure 6:**
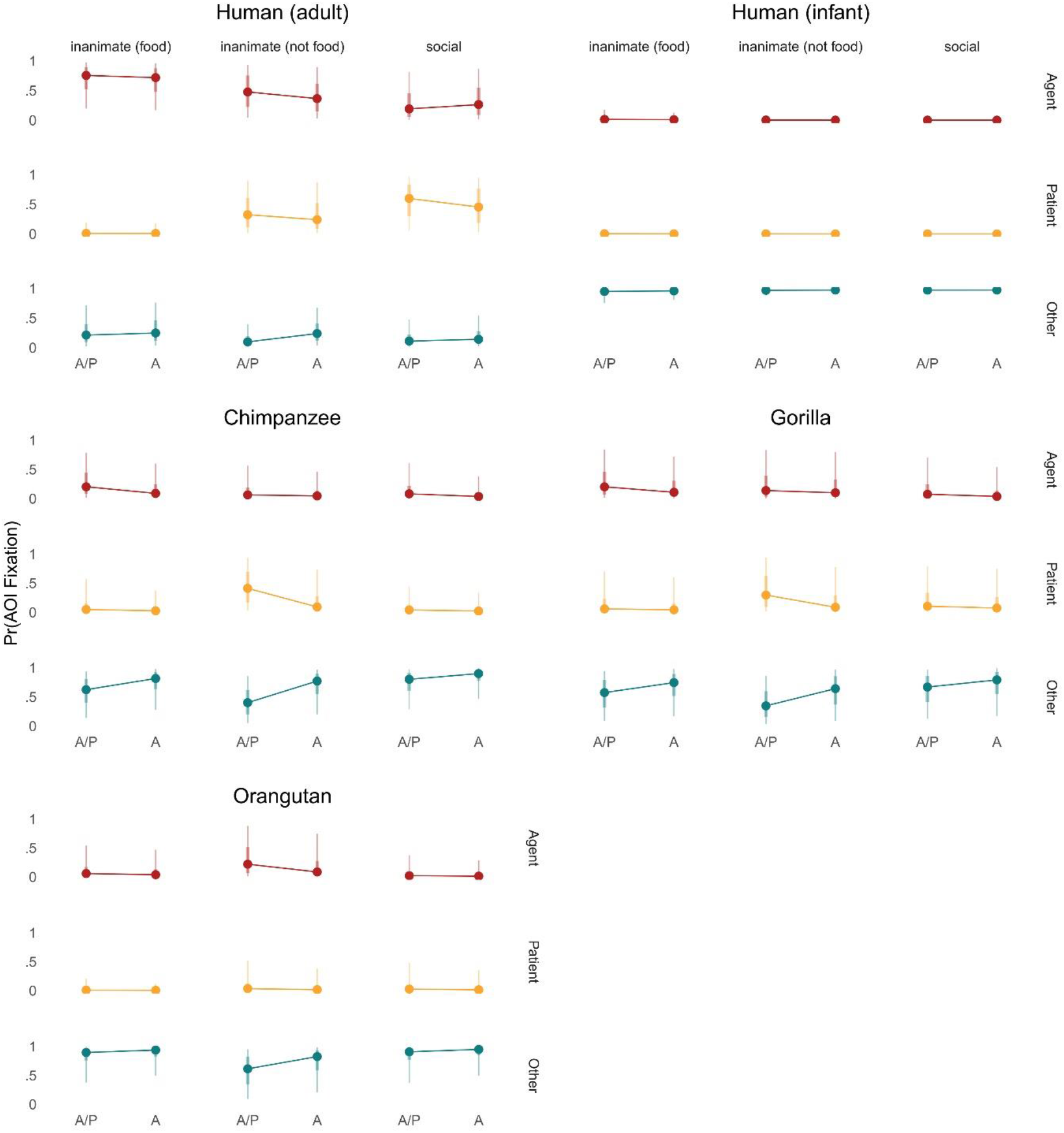
AOI fixation probability by movement difference between agent and patient. A = agent moves more than patient. A/P = agent and patient move a similar amount. Colors depict the agent (red), patient (orange), and other information (turquoise).

## Notes

### Competing Interest Statement

The authors have declared no competing interest.

